# Decoupling Transcription from Cell Fate Preserves Germ Cell Totipotency

**DOI:** 10.64898/2026.09.17.752471

**Authors:** Sherilyn Grill, Arjuna Rajakumar, Anaïs Tsai, Claire Martel, Lake V. Palmeri, Xinlei Gao, Amelie A. Raz, Ruth Lehmann

## Abstract

Only germ cells can give rise to a totipotent embryo, yet a conserved transcriptional program for germ cell identity remains elusive. Profiling *Drosophila* primordial germ cells (PGCs) across embryogenesis, we find that nascent PGCs are not silent, as previously thought, but instead transcribe the same totipotency program as the early embryo, reflecting an open chromatin state. This chromatin state, established by the pioneer factor Zelda, persists in PGCs throughout development. The associated transcriptional program normally drives somatic differentiation, but in PGCs the conserved RNA-regulator Nanos represses its translation, leaving germline chromatin uncommitted. Cells that fail to establish the totipotent state die before reaching the gonad. Thus, germ cells preserve developmental potential not by restricting transcription, but by preventing transcripts from determining cell fate.

## Main text

Germ cells are the sole link between generations and, upon fertilization, are uniquely capable of generating every cell type in the body. Yet no dedicated transcriptional program for primordial germ cell (PGC) fate has been identified, leaving the molecular basis of germline specification unresolved (*1*). Set aside early in embryogenesis, primordial germ cells are segregated from somatic lineages, each committed to distinct developmental fates (*2*). Cell fate is normally established through a stepwise process, in which a cell transcribes a lineage-specific program that drives its differentiation, progressively restricting the chromatin until the differentiated state becomes irreversible (*3, 4*). Somatic lineages are thus defined by dedicated master-regulator transcription factors that activate these programs. Germ cell identity, however, is characterized instead by global transcriptional repression (*5–8*). Consistent with this, the maternal-to-zygotic transition is delayed in the germline, with maternal transcripts retained longer and zygotic genome activation occurring later than in somatic lineages (*9, 10*). How germ cells establish and maintain a distinct identity in the absence of a defining transcriptional program remains a fundamental unsolved question in developmental biology.

In *Drosophila*, the germline is established when syncytial nuclei migrate to the posterior pole of the embryo and bud off to form pole cells, the first cells of the embryo and the precursors of the primordial germ cells (*6*). Across animals, roughly 30% of primordial germ cells are lost before reaching the gonad, yet the basis of this conserved attrition is unknown (*11–15*). Whether all of these cells have the capacity to become functional PGCs has not been resolved in any organism. The lost cells may simply be normal germ cells that are eliminated, or they may be cells that fail to complete a discrete molecular transition required for germ cell fate, and are lost as a result. Addressing this requires capturing the earliest molecular events of germ cell specification as they unfold. These events occur during the brief transition from the maternal to the zygotic transcriptome, a window that is difficult to capture at sufficient resolution in most organisms. *Drosophila* offers a unique advantage: collections of timed embryos enable near-continuous temporal sampling, allowing us to capture the precise sequence of molecular events during germ cell specification at single-cell resolution, with minutes rather than hours separating developmental stages. To capture the transcriptional events of germ cell specification at this resolution, we generated a comprehensive single-cell transcriptomic atlas of *Drosophila* primordial germ cells spanning embryonic development.

We collected embryos spanning pole cell formation through gonad coalescence (1.5–12 hours post-fertilization (hpf)) (Fig. 1A, fig. S1A). We defined time windows to capture germ cells across four developmental stages: cellularization, specification, migration, and gonad coalescence (Fig. 1A). For each developmental window, embryos were dissociated and germ cells were isolated by FACS using a GFP transgene driven by the Vasa promoter and a live-cell viability marker (fig. S1B). Two biological replicates per time window were processed independently for single-cell RNA sequencing. Data were analyzed using Seurat for clustering and Monocle3 for pseudotime analysis (see Methods).

**Figure 1:**
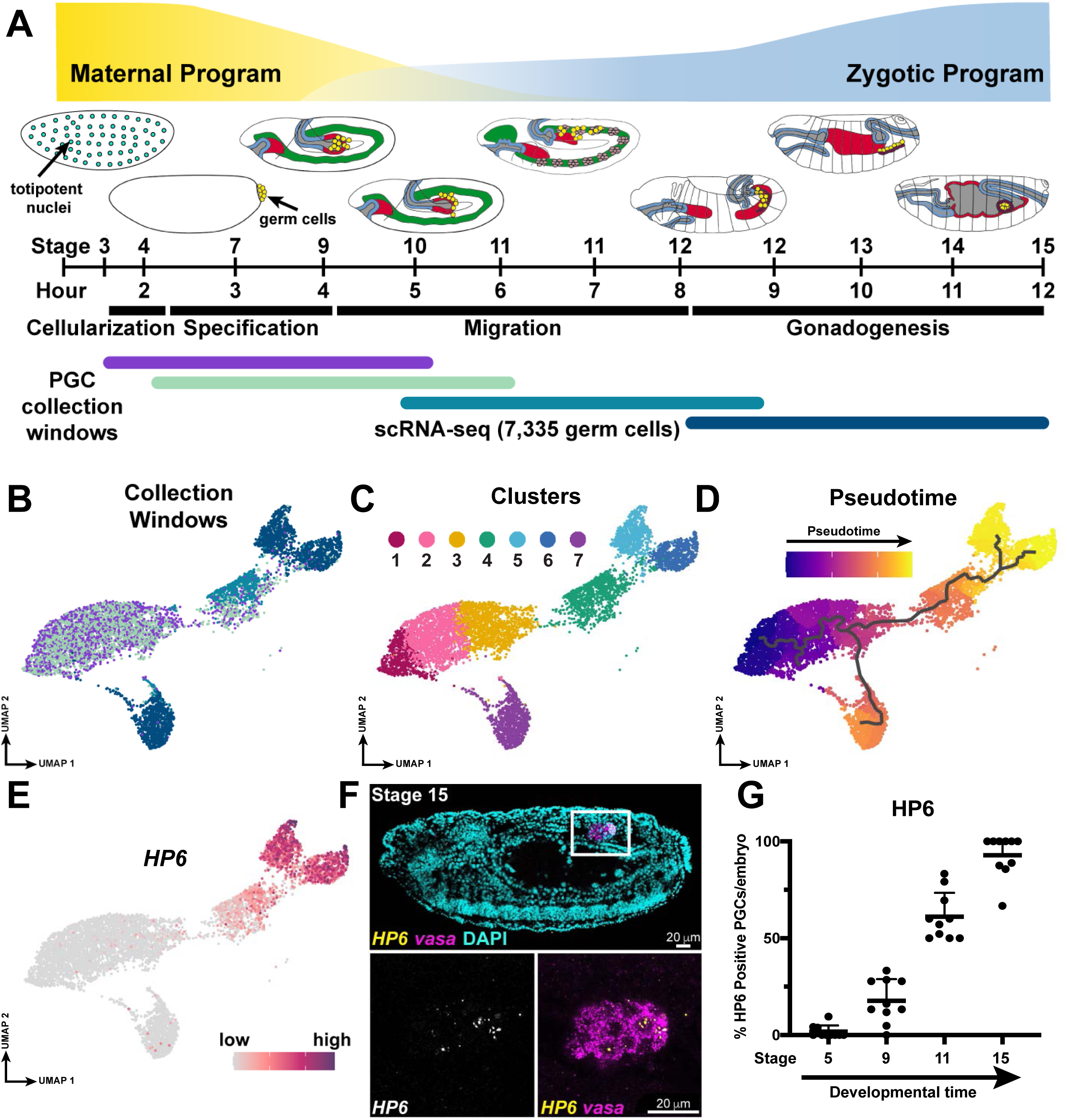
Atlas of germ cell development across embryogenesis. A. Schematic of the maternal-to-zygotic transition in primordial germ cells. Maternally deposited transcripts (yellow) are progressively degraded while zygotic transcription (blue) is gradually activated. Embryo schematics are adapted from (*68*), with tissues color-coded: cyan, totipotent nuclei; yellow, germ cells; green, mesoderm; blue, foregut/hindgut; red, midgut; light purple, lateral mesoderm; dark purple, somatic gonadal precursors. Developmental stages and hours post fertilization are indicated. Black bars mark key germ cell developmental events. The four overlapping embryo collection windows spanning primordial germ cell development are shown below. Germ cells from P{vasa-GFP} embryos were collected, sorted, and used for single-cell RNA sequencing. B. UMAP visualization of scRNA-seq data from isolated germ cells across four developmental windows, colored by timepoint. About 5% of embryos in these early collections (purple and light green) were contaminated with later-stage embryos due to egg retention in females. These ‘older’ germ cells cluster according to their developmental age. C. UMAP colored by cluster identity, revealing transcriptionally distinct populations within and across developmental stages. D. Pseudotime analysis of the germ cell population, showing a developmental trajectory progressing from early to late embryogenesis. E. Feature plot showing expression of *HP6*, a representative gene that is expressed during the major wave of ZGA in the germline. F. *In vivo* validation of *HP6* dynamics by mRNA *in situ* hybridization using HCR (hybridization chain reaction). *HP6* transcript in yellow, germ cells marked by *vasa* transcript in magenta, nuclei shown with DAPI. Full embryo (top) with high-magnification inset of germ cells (bottom); scale bars, 20 µm. G. Quantification of HCR-stained embryos recapitulates the transcriptional dynamics observed by scRNA-seq. Each dot represents one embryo.

Post-processing quality control identified small numbers of contaminating somatic cells, including somatic gonadal precursors, consistent with their known low-level vasa expression (*16*). The germ cell population was cleanly separable across replicates. Dimensionality reduction revealed a single branching structure spanning the full course of embryonic germline development from pole cell formation to 12 hpf. The resulting UMAP resembled a whale, with the earliest timepoints occupying the nose and later timepoints represented in the body and tail (Fig. 1B). Clustering analysis confirmed this developmental trajectory based on the expression dynamics of known maternal and zygotic markers (Fig. 1C, E, fig. S1C-M) (*17–20*). Pseudotime analysis identified a single primary developmental trajectory from the earliest (nose) to the latest (tail) clusters, indicating that germ cells progress through one continuous developmental path throughout embryogenesis (Fig. 1D).

### The germline maternal-to-zygotic transition at single-cell resolution

By sampling germ cell development at near-continuous temporal resolution, our atlas captures the germline maternal-to-zygotic transition as a continuous process. This resolution allows transient and rapidly changing transcripts to be detected as they unfold, rather than averaged across broad developmental windows. At the earliest timepoints, the germ cell transcriptome is dominated by maternally deposited RNAs. *Oskar* (*osk*), which is rapidly degraded upon pole cell cellularization, marks the most nascent pole cells at the earliest pseudotime in the nose of the whale, serving as a molecular benchmark of germ cell age (fig. S1F) (*21*). *Pgc*, a representative maternal transcript, is detected in all early germ cells in our transcriptomic dataset and is degraded over developmental time (fig. S1C). HCR *in situ* hybridization confirms this pattern *in vivo*, with *pgc* RNA ubiquitous in early pole cells and largely absent by 7 hours post fertilization (fig. S1G, H).

Between the early and middle timepoints, maternal transcripts are degraded, and the minor wave of zygotic genome activation (ZGA) begins in the germline. Consistent with its zygotic activation in germ cells at 3-5 hours of development (*9*), *zen* is expressed during the minor wave of ZGA in our dataset (fig. S1D). HCR *in situ* hybridization confirms this pattern *in vivo*, with robust *zen*transcription by stage 9 (fig. S1I, J). Because our transcriptomic data cannot distinguish maternally deposited from zygotically transcribed *vasa*, we used HCR probes targeting intronic sequences to specifically visualize nascent *vasa* transcription. Consistent with prior reports, zygotic *vasa* expression also initiates during the minor wave of ZGA in the germline at stage 9, and persists through gonad coalescence (fig. S1E, K, L) (*22*). More broadly, transcripts with both maternal and zygotic contributions appear uniformly expressed across all cells in our dataset, precluding identification of zygotic transcription onset for this class of genes (fig. S1E). Robust genome activation occurs during the major wave of ZGA as germ cells reach the somatic gonad, corresponding to the tail of the whale. *Heterochromatin protein 6* (*HP6*), a representative gene expressed during the major wave of ZGA, peaks at stage 15 (Fig. 1E-G) (*19*).

Intriguingly, the tail of the whale bifurcates into two clusters corresponding to XX and XY PGCs, as evidenced by the exclusive expression of the Y-linked gene *FDY* in one cluster (fig. S1M). Differential gene expression between these clusters is driven primarily by the 2-fold higher expression of X-linked genes in XX germ cells, reflecting the absence of dosage compensation at this stage of germ cell development (*23, 24*). Consistent with this, 99 of the 100 most significant markers distinguishing the two clusters are X-linked; the sole exception is *FDY*, a Y-linked gene expressed specifically in the XY population (table S1, fig. S1M). The ability to resolve this 2-fold difference underscores the sensitivity of our atlas. Prior studies have characterized the *Drosophila* primordial germ cell transcriptome at bulk and single-cell resolution (*9, 23, 25*). Building on these analyses, our atlas captures the rapid yet coordinated transition from maternal genome to germline program with greater temporal resolution and coverage, revealing regulatory dynamics not previously resolved.

### PGCs initiate zygotic transcription immediately after formation

For over three decades, pole cells have been described as transcriptionally repressed upon formation, with the first germline zygotic transcripts identified during gastrulation (*8, 9, 22, 26, 27*). Prior studies established that germline ZGA occurs in two waves following this initial period of transcriptional quiescence (*9, 22*). We set out to resolve these dynamics at single-cell resolution. To do so, we identified all transcripts that change in abundance across developmental time, excluding constitutively expressed genes, and grouped them by their expression dynamics (Fig. 2A). Unexpectedly, this revealed that newly formed pole cells are not transcriptionally quiescent. Instead, a discrete set of approximately 30 genes is activated immediately following pole cell cellularization, prior to any previously described wave of ZGA (Fig. 2A, fig. S2A, B, table S2) (*9*). Together, this analysis defined four transcriptional classes based on our *in vivo* validation and expression dynamics across pseudotime: Class I, maternally deposited transcripts that are progressively degraded; Class II, the newly identified genes that are transiently activated immediately after pole cell cellularization; Class III, activated during the minor wave of ZGA; and Class IV, activated during the major wave of ZGA (Fig. 2A, fig. S2A,B). Across pseudotime, Class II genes share a coordinated, transient expression profile that peaks during the germ cell specification window, anti-correlated with the decline of the maternal transcriptome and preceding both known waves of ZGA (fig. S2A, B). The one exception is the Class II gene *Taf12L*, which is expressed more continuously throughout germline development (fig. S2B). Together, Class II expression marks the transition from the maternal program to a specified germ cell transcriptome. This early transcription likely escaped prior detection because it represents only a small fraction of the pole cell transcriptome: 0.14% of transcripts in pole cells, which are otherwise dominated by maternal mRNAs. We confirmed Class II transcription *in vivo* using HCR *in situ* hybridization. *Amos*, a representative Class II gene, is absent from the germ plasm before pole cell cellularization, appears in pole cells, peaks at stage 5, and becomes undetectable in migrating PGCs by stage 9 (Fig. 2B-D, for developmental staging see Fig. 1A). This early zygotic expression was detected across every Class II gene examined, including *amos*, *sisA*, *ato*, *Elba1*, and *Taf12L*, which are all expressed in pole cells beginning at stage 4 (Fig. 2B-E, fig. S2B-F). Notably, Class II expression is stochastic in the germline: at peak expression, some pole cells transcribe Class II genes robustly, others weakly, and others not at all, with no apparent positional bias within the germ cell cluster (Fig. 2B-E, fig. S2C-F).

**Figure 2:**
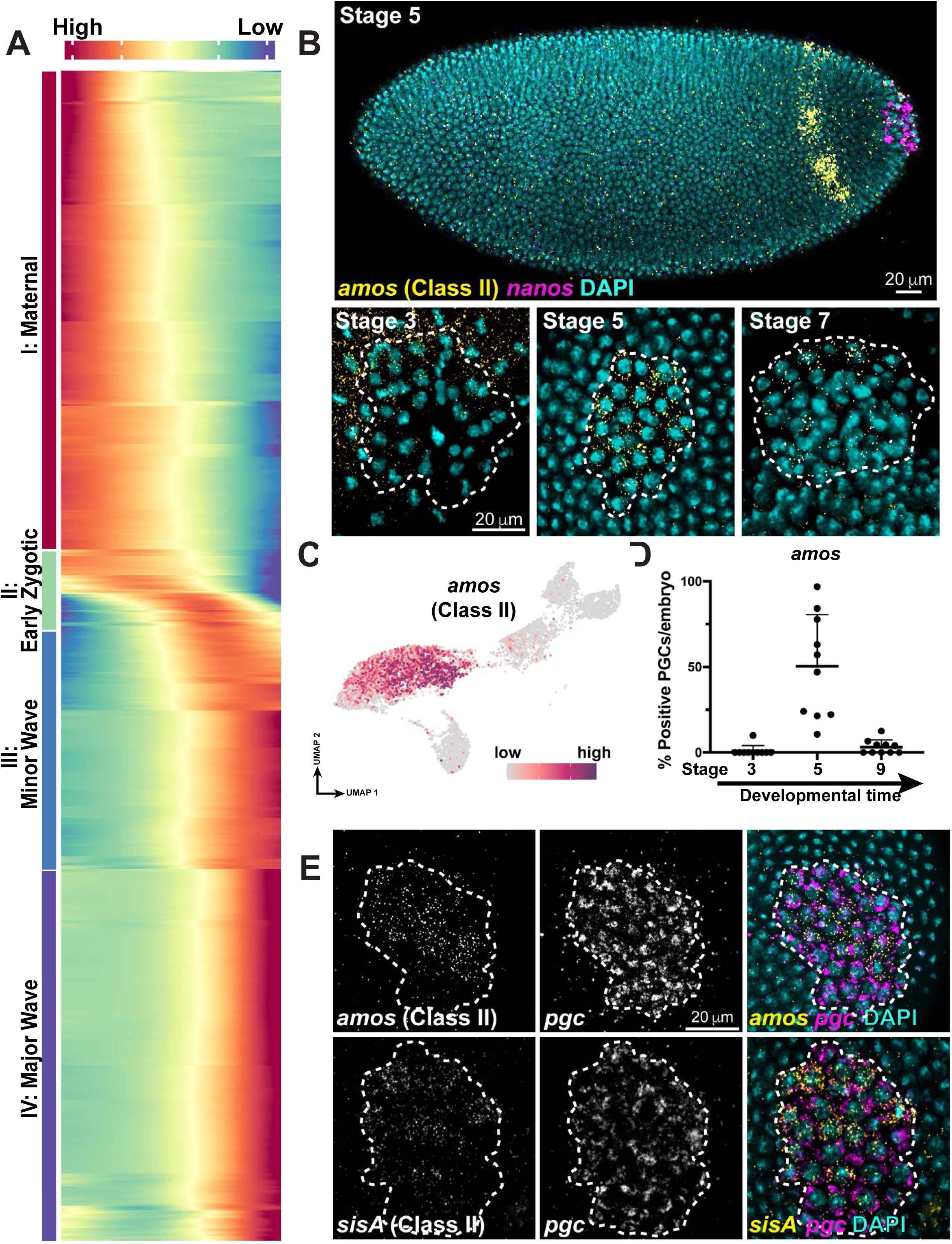
Nascent germ cells activate a discrete early zygotic transcriptional program at their formation. A. Heatmap of genes that significantly change in expression across pseudotime, revealing four classes: Class I (maternally deposited), Class II (early zygotic), Class III (minor ZGA), and Class IV (major ZGA). B. HCR *in situ* hybridization of *amos* transcript (yellow) in germ cells marked by *nanos* (magenta). Top: stage 5 embryo, with *amos* detected in pole cells and a posterior somatic stripe. Bottom: posterior-mounted embryos showing *amos* expression in the germline (dotted outline) beginning at stage 3, peaking at stage 5, and declining by stage 7. Nuclei in DAPI (cyan). Scale bars, 20 µm. C. Feature plot showing expression of a representative Class II gene, *amos*, from scRNA-seq analysis. D. Quantification of *amos*-positive PGCs across developmental stages in HCR-stained embryos. Each dot represents one embryo. E. HCR *in situ* hybridization of posterior-mounted wild-type embryos. Pole cells shown inside the dotted outline are marked by *pgc* transcript (magenta) and express *amos* (yellow, top) or *sisA* (yellow, bottom). Scale bar, 20 µm.

Together, these findings challenge the long-standing view that pole cells are transcriptionally silent. Instead, nascent pole cells activate a discrete transcriptional program at the very moment of germ cell specification, defining a distinct molecular event that has, until now, gone undetected.

### Germ cells co-opt the founding transcriptional program of the totipotent blastomeres

After identifying a discrete class of transiently expressed genes in pole cells, we next asked whether these genes encode germline-specific factors that might act as master regulators of germ cell fate. Surprisingly, we found that Class II genes are not germline determinants, but instead are nearly identical to the first genes transcribed in the *Drosophila* embryo (Fig. 3A, fig. S3A, Table S2) (*28–30*). Like totipotent mammalian blastomeres, the *Drosophila* blastoderm nuclei are competent to generate all cell types in the organism, and in both systems this totipotent state is rapidly lost as cells commit to distinct lineages (*31, 32*). We therefore refer to the syncytial blastoderm nuclei in *Drosophila* as totipotent blastomeres, drawing an explicit parallel to their mammalian counterparts. Our results suggest that the first transcriptional program of Drosophila germ cells resembles the totipotent transcriptional state of the embryonic blastomeres.

**Figure 3:**
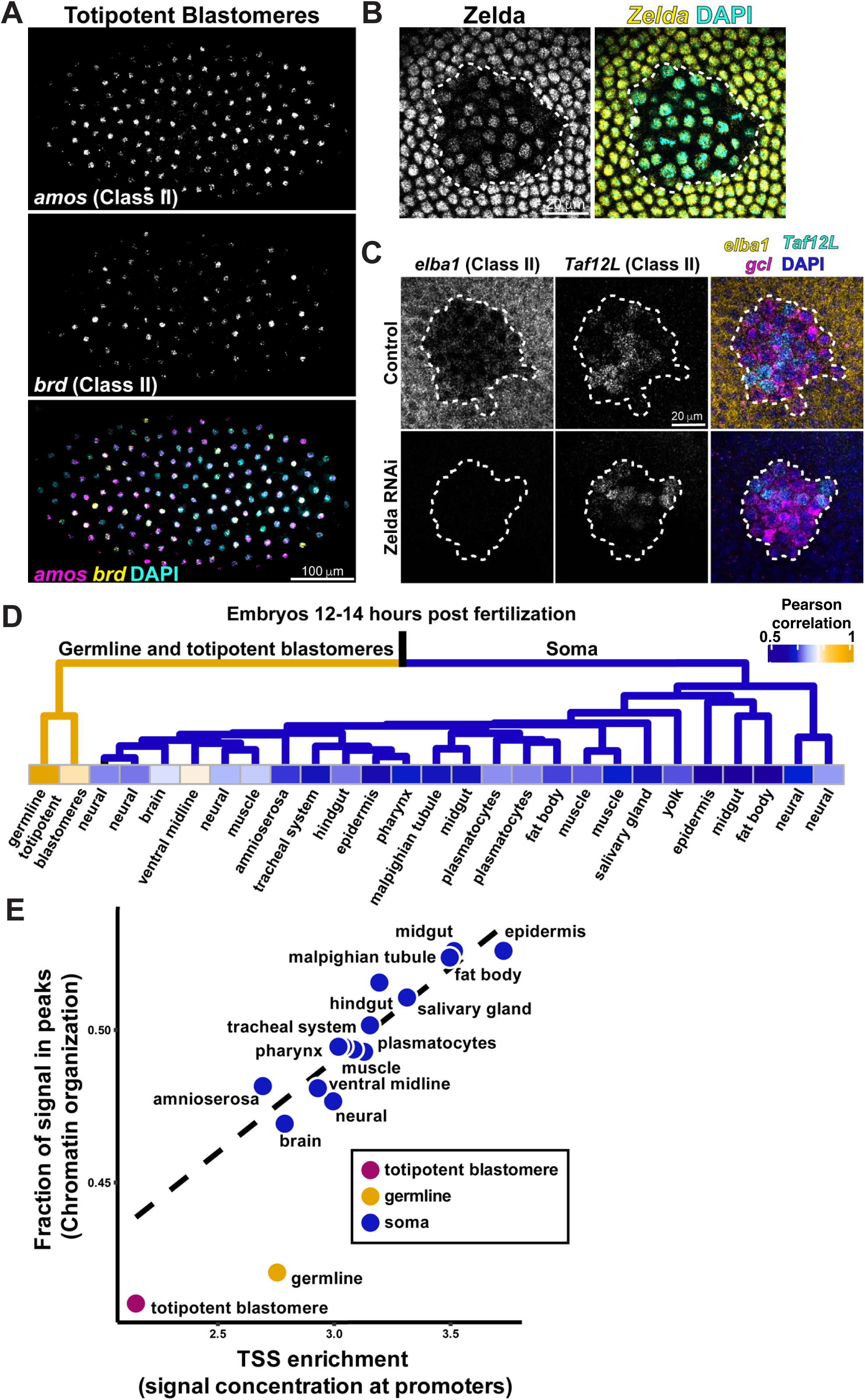
Germ cells maintain a totipotent identity throughout embryogenesis. A. HCR *in situ* hybridization of a laterally mounted stage 3 embryo showing the totipotent blastomeres. *Amos* (magenta) and *brd* (yellow), two representative Class II genes, are expressed in the blastomeres. Nuclei shown with DAPI (cyan). Scale bar, 100 µm. B. Zelda protein is present in stage 4 pole cell nuclei. Immunofluorescence staining of Zelda protein (yellow) in pole cell and somatic nuclei, with higher levels in the soma than in the pole cells. Pole cells are outlined (dotted line). Nuclei shown with DAPI (cyan). Quantification shown in Fig. 4D. Scale bar, 20 µm. C. Zelda is required for Class II gene expression. HCR *in situ* hybridization of control and Zelda knockdown stage 4 embryos showing *elba1* (yellow) and *Taf12L* (cyan), two Class II genes. Germ cells are marked by *gcl* (magenta) and outlined (dotted line). *elba1* expression is lost upon Zelda knockdown in both the pole cells and the soma, while *Taf12L* expression is maintained. Scale bar, 20 µm. D. Heatmap of pairwise Pearson correlation coefficients across snATAC-seq chromatin accessibility profiles of all identified cell types in embryos 12-14 hours post fertilization (hpf) together with totipotent blastomeres from embryos 0-2 hpf and compared to germ cells from embryos 12-14 hpf. The dendrogram (top) shows that PGCs from embryos 12-14 hpf cluster with the totipotent blastomeres (yellow) rather than with differentiated somatic cell types (blue), indicating that germ cells retain a totipotent-like chromatin state throughout embryogenesis. snATAC-seq data from (*41*). E. Scatter plot of chromatin openness metrics for each cell type at 12-14 hpf, plotting TSS enrichment (signal concentration at promoters) against the fraction of reads in peaks, a measure of chromatin organization. A linear fit to the somatic cell types is shown (dashed line). Germ cells fall well below the somatic trend, displaying a somatic-like TSS enrichment but a markedly lower fraction of reads in peaks, indicating that germ cell chromatin remains globally unorganized with less peak enrichment despite their differentiated state. snATAC-seq data from (*41*).

We find that 35 of the 36 Class II genes transcribed during PGC specification are also transcribed in totipotent blastomeres (Table S2, Fig. 3A, fig. S3A) (*28–30, 33*). The one exception is *Taf12L,* which is expressed exclusively in the germline (fig. S3A). In totipotent blastomeres, the pioneer transcription factor Zelda activates Class II genes by opening chromatin and driving the earliest wave of ZGA, and many of the Class II genes carry among the highest Zelda ChIP-seq signal in the genome (Table S2) (*33–35*). Because germ cells transcribe these same genes, we asked whether Zelda also drives their expression in the germline.

We find that Zelda protein is present in pole cell nuclei (Fig. 3B). To test whether Zelda is required for Class II transcription in the germline, we performed maternal Zelda knockdown. Class II gene expression is abolished in both the somatic cells and pole cells upon loss of maternal Zelda (Fig. 3C, Table S2), demonstrating that the same pioneer factor drives early transcription in both cell types. Notably, *Taf12L* is activated independently of Zelda in pole cells, distinguishing it from the broader Class II program (Fig. 3C).

Strikingly, many of these Class II genes encode somatic transcription factors that drive cell fate specification in somatic, non-germline lineages, including the proneural transcription factors *amos* and *ato* (Table S2) (*36–38*). These genes are characterized by their somatic loss-of-function phenotypes and have no reported role in germline development, consistent with Class II genes representing a shared totipotency program rather than specific germline determinants. *Taf12L*, the sole germline-specific Class II gene, is the only member with a known germline function in adult spermatogenesis (*39, 40*). We therefore directly tested its requirement in germ cell development by CRISPR knockout, and found no detectable effect on PGC number (fig. S3B). Together, these findings suggest that germ cell specification is not driven by a unique transcriptional program. Instead, germ cells re-activate the embryo’s own founding totipotency program in a new cellular context. This program does not, however, contain obvious determinants of germline identity. Many Class II genes encode somatic master regulators, and in the totipotent blastomeres Class II genes are often transcribed as short, abortive products rather than functional mRNAs (*29*). The shared expression of these genes therefore reflects a common transcriptional state between pole cells and totipotent blastomeres, but not one that determines germline fate.

### Class II genes escape Pgc-mediated repression

The activation of a discrete transcriptional program in pole cells was unexpected, as pole cells were thought to be completely transcriptionally silent. This silence is maintained by the maternally deposited protein Polar Granule Component (Pgc), which blocks the release of paused RNAPII into productive elongation by inhibiting Serine 2 phosphorylation in the CTD (*17, 18, 26*). We therefore asked how Class II genes escape this repression.

We found that Class II genes are transcribed in pole cells that contain *pgc* RNA (Fig. 2E, fig. S2E). These same pole cells lack RNAPIIS2p, the mark of productively elongating polymerase, despite its strong presence in the surrounding soma (fig. S2D, E), consistent with Pgc-mediated repression being active. H3K4me3, an activating histone mark associated with productive transcription, is likewise high in the soma but absent in Class II-expressing pole cells, suggesting transcription of Class II genes occurs in the absence of productive elongation. (fig. S2F). Yet many of the resulting Class II transcripts are complete: they are full-length and polyadenylated, detected in our poly(A) capture scRNA-seq atlas and identified by HCR in the cytoplasm. We find that Class II genes are characteristically short and largely intronless, and in the totipotent blastomeres they are transcribed without promoter-proximal pausing (Table S2) (*28–30*). Because these genes do not pause, they do not require pause release, the very step that Pgc inhibits. The same property that allows Class II genes to be transcribed rapidly in the blastomeres therefore renders them insensitive to Pgc repression in the germline.

### Germ cells maintain a totipotent chromatin state throughout embryogenesis

Class II genes are not established determinants of germline fate. Their shared transcription in pole cells and totipotent blastomeres is therefore unlikely to reflect a common regulatory program, and may instead be a consequence of a shared chromatin state that allows the same genes to be expressed in both cell types. We asked whether germ cells and totipotent blastomeres share a common chromatin landscape.

We compared chromatin accessibility profiles across germ cells, totipotent blastomeres, and all major somatic cell types throughout embryonic development using published single nucleus ATAC-sequencing (snATAC-seq) data (*41*). As totipotent blastomeres are present only in the early embryo, we used the chromatin of 0-2 hour blastomeres as the totipotent reference, comparing germ cells and somatic lineages at all later stages against it. In 2-4 hour embryos, nascent germ cells, somatic lineages, and 0-2 hour blastomeres display broadly similar chromatin landscapes, reflecting the undifferentiated state of the early embryo (fig. S3C). As development proceeds, somatic lineages progressively organize their chromatin and commit to their differentiated fates. Strikingly, germ cells do not follow this trajectory. Instead, they maintain a chromatin landscape that more closely resembles that of the totipotent blastomeres than any differentiated somatic lineage (Fig. 3D, fig. S3D). At 12-14 hours, long after somatic nuclei have committed to their fates and restricted their chromatin, PGCs that have migrated and coalesced with the gonad still retain an unrestricted chromatin state characteristic of the totipotent blastomeres (Fig. 3D, fig. S3D). Pairwise correlation analysis confirms that germ cells cluster with the 0-2 hour totipotent blastomeres rather than with any differentiating somatic cell type, and dendrogram analysis places 12-14 hour germ cells and 0-2 hour totipotent blastomeres in a distinct clade from all other 12-14 hour somatic lineages (Fig. 3D, fig. S3D).

Pluripotent and totipotent cells are characterized by globally open chromatin, a transcriptionally permissive state associated with developmental plasticity, whereas differentiation drives chromatin compaction and concentrates accessibility at defined regulatory elements (*42, 43*). To assess where germ cells fall along this spectrum, we examined two features of chromatin organization: enrichment of accessibility at transcription start sites (TSS) and the fraction of accessible signal concentrated within peaks (FSiP). FSiP is a measure of genome organization, indicating whether accessibility is concentrated at discrete sites or spread broadly across the genome. Germ cells show TSS enrichment comparable to differentiated somatic lineages, consistent with their status as a specified cell type. Yet their fraction of signal in peaks is the lowest of any cell type in the 12-14 hour embryo (Fig. 3E), indicating that germ cell accessibility is broadly distributed across the genome rather than concentrated at discrete regulatory elements. As somatic cells differentiate, they restrict their chromatin, losing the broad accessibility of the early embryo and concentrating accessibility at the promoters of their specific program (Fig. 3E). Germ cells instead retain the broad accessibility of the early embryo and acquire promoter accessibility on top of it, maintaining an unrestricted chromatin landscape (Fig. 3E). By combining an organized promoter landscape while preserving the broadly accessible chromatin of the early embryo, germ cells may reconcile their differentiated identity with their unique capacity to regenerate a totipotent embryo upon fertilization.

We propose that this unrestricted chromatin state is a defining feature of germ cell specification. The transcription of Class II genes is likely a consequence of this permissive chromatin combined with their ability to escape Pgc-mediated repression. As a result, germ cells are left poised to differentiate inappropriately.

### Nanos represses translation in the germline to protect germ cell identity

We find that germ cells maintain a totipotent chromatin state that permits transcription of a broad set of somatic fate-specifying genes (Table S2). Germline and somatic fates are fundamentally opposed, and the germline is defined by its exclusion from somatic differentiation. Yet at the level of transcription, germ cells resemble the very somatic cells they must not become. Because many Class II genes encode somatic fate-specifying transcription factors, we asked whether their transcription in the germline produces functional protein. To address this, we generated an endogenously GFP-tagged *amos* knock-in line, which faithfully recapitulates endogenous *amos* expression dynamics in pole cells, drives GFP expression in neurons, and produces maternally deposited Amos-GFP protein in early embryos, confirming its functionality (fig. S4A-C). Despite active *amos* transcription in pole cells, Amos protein is largely absent from the germline (Fig. 4A, B). The decoupling of *amos* transcription from Amos protein accumulation in pole cells indicates that Class II transcripts are subject to translational repression.

**Figure 4:**
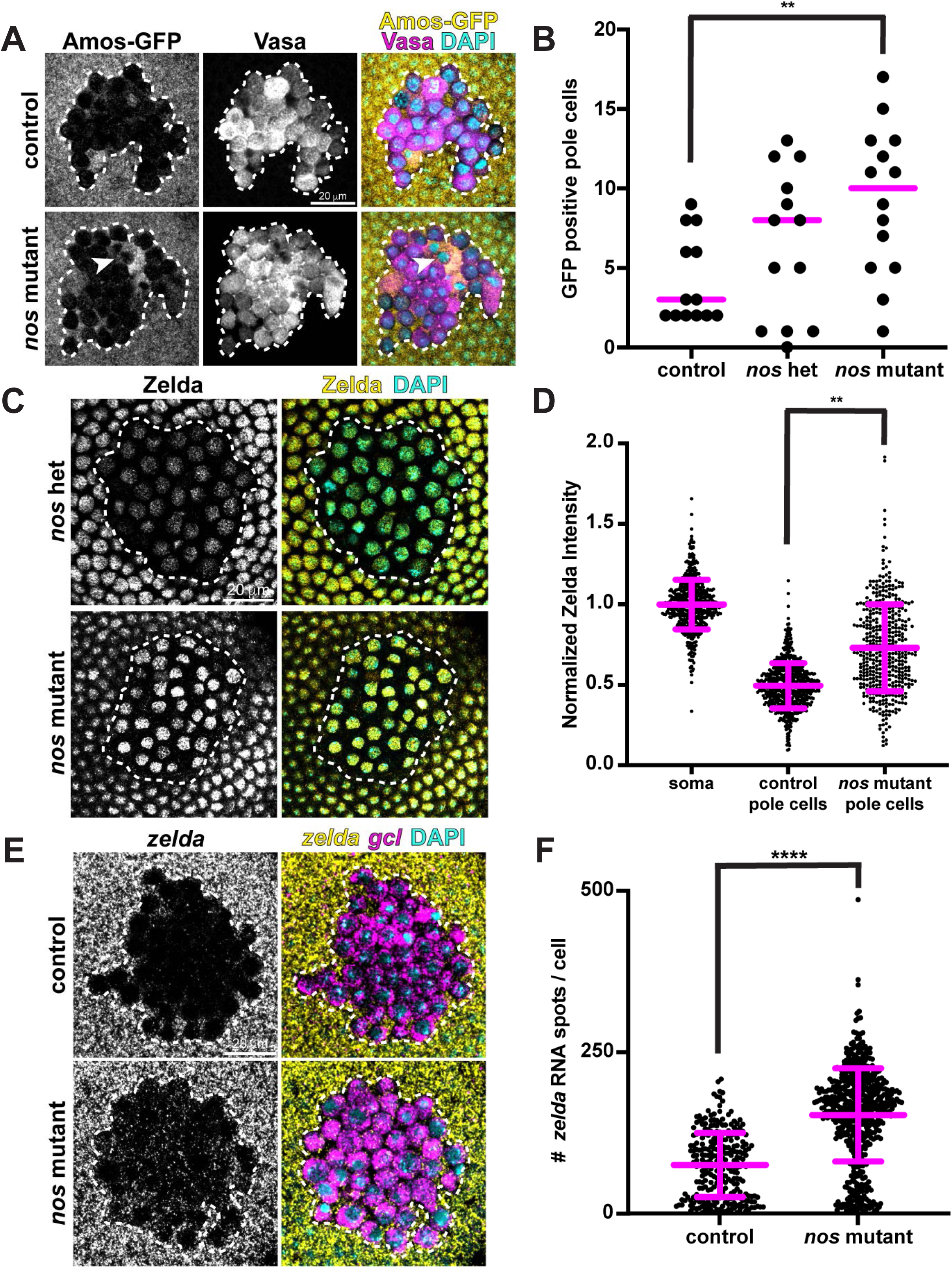
Nanos prevents class II translation in pole cells. A. HCR Immunofluorescence staining of sibling control and *nanos* mutant embryos showing Amos-GFP (yellow) and Vasa protein (magenta). Dotted lines encompass pole cells. Arrowheads mark Amos-GFP-positive pole cells. Nuclei shown with DAPI (cyan). Scale bar, 20 µm. Wild-type pole cells do not accumulate significant Amos protein, while *nanos* mutant pole cells aberrantly accumulate Amos protein. B. Quantification of the number of GFP-positive pole cells per embryo from embryos stained as in (A). Sibling control, heterozygous, and *nanos* mutant embryos were stained and quantified. Each dot represents one embryo; bars indicate mean ± SD. Comparison between sibling-matched control embryos and *nanos* mutant embryos was performed using an unpaired Welch’s t-test (n = 13 and 14 embryos, respectively); **P = 0.0024. C. Nanos prevents Zelda accumulation in pole cells. Immunofluorescence staining of control heterozygous and *nanos* mutant embryos showing Zelda protein (yellow) and DNA (cyan) in pole cell and somatic nuclei. Pole cells outlined (dotted line). Zelda protein is elevated in *nanos* mutant pole cells relative to controls, with some pole cells reaching the same fluorescence intensity as the soma. Scale bar, 20 µm. D. Quantification of Zelda protein levels in soma, control pole cells, and *nanos* mutant pole cells from embryos stained as in (C). Each dot represents one nucleus; bars indicate mean ± SD. Comparison between control and *nanos* mutant PGCs was performed on per-embryo means using a Wilcoxon rank-sum test (n= 23 and 16 embryos, respectively); **P = 0.0044. E. *nanos* mutant pole cells retain *zelda* transcript. HCR *in situ* hybridization of control and *nanos* mutant embryos showing *zelda* (yellow) and *gcl* (magenta, germ cell marker). Pole cells outlined (dotted line). Scale bar, 20 µm. F. Quantification of *zelda* mRNA signal in pole cells from embryos stained as in (E). Each dot represents one cell; bars indicate mean ± SD. Comparison between control and *nanos* mutant pole cells was performed using an unpaired Welch’s t-test (n = 273 and 561 pole cells, from 8 and 14 embryos, respectively); ****P < 0.0001.

Translational control has long been considered a hallmark mechanism of germ cell regulation, making it a likely regulator of Class II repression in the germline. Specifically, the RNA-binding protein Nanos is a conserved germline determinant required for germline identity across animals (*44–46*). Nanos mRNA is deposited maternally into the embryo, locally enriched and translated in the germ plasm, where it promotes mRNA degradation and represses translation (*47, 48*). We thus tested whether Nanos represses Class II translation. Amos protein, which is absent from wild-type pole cells despite active *amos* transcription, accumulates significantly in the pole cells of embryos from *nanos* mutant females (hereafter referred to as *nanos* mutant) (Fig. 4A, B). Consistent with Nanos repressing the Class II program, 23 Class II genes are significant targets of Nanos, with 30 showing a trend toward Nanos-mediated depletion, in a genome-wide analysis (Table S2) (*49*). Nanos is therefore required to limit Class II protein accumulation in the germline.

Nanos also limits Class II mRNA abundance. Transcript levels of multiple Class II genes, including *elba1* and *sisA*, were elevated in maternal *nanos* mutant pole cells (fig. S5A, B). The coordinated increase across many Class II genes suggested that *zelda*, their shared activator, is an upstream target of Nanos. Consistent with this, Zelda protein was elevated in *nanos* mutant germ cells (Fig. 4C, D). Examining *zelda* RNA directly, we found that at nuclear cycle 11 (∼1.5 hours post-fertilization), *zelda* RNA is present in both wild-type and *nanos* mutant pole cells at levels similar to the surrounding soma (fig. S5C). By nuclear cycle 14 (∼3 hours post-fertilization), *zelda* RNA was cleared from control pole cells but persisted in *nanos* mutant pole cells (Fig. 4E, F). We propose that in the absence of Nanos, *zelda* RNA escapes degradation and is translated, producing elevated Zelda protein that in turn drives increased Class II transcription in the germline. Consistent with its Zelda-independent activation, *Taf12L* transcription remained unchanged in *nanos* mutant pole cells (fig. S5B). This increased Class II transcription in *nanos* mutants was not due to changes in the RNAPII machinery. RNAPIIS5p and RNAPIIS2p levels are unchanged in *nanos* mutant germ cells relative to controls (fig. S5D-G), indicating that the elevated Class II transcription reflects increased Zelda activity rather than direct modulation of RNAPII machinery.

Together, these data resolve a fundamental paradox. In the totipotent blastomeres, the founding transcriptional program initiates somatic differentiation. Germ cells activate this same initial program, yet Nanos blocks its downstream consequences by repressing translation of somatic determinants and limiting Zelda to prevent differentiation. Nanos is present in the germline throughout development, providing continuous protection. As a result, germ cell chromatin stays uncommitted and totipotent, while germline fate is set at the level of translation. In the germline, translational control decouples transcription from cell fate specification.

### A transcriptionally stalled population of PGCs fails specification and dies: the lost children

If activation of the totipotent program defines germ cell specification, then failure to activate it should compromise germ cell fate. Indeed, we find that not all pole cells activate this program: a subset fails specification entirely and dies before reaching the gonad, indicating that activation of the totipotency program is essential for primordial germ cell fate.

Among the germ cells in our dataset, we noticed a striking outlier: a cluster of late-stage germ cells with a transcriptome resembling newly formed pole cells rather than that of their developmental age (Fig. 1B, C; cluster 7-purple). This cluster was isolated at the latest developmental timepoint, yet these cells surprisingly contain a transcriptome composed entirely of maternal transcripts, with no detectable zygotic transcription (Fig. 1E, Fig. 2C, fig. S1C, D, M). Although these cells retained their maternal transcriptome far longer than healthy germ cells, maternal transcript levels in this cluster were modestly lower than in newly formed pole cells, indicating that some maternal transcript loss occurs over time (fig. S1C). Using computational tools that correct for technical variation between samples, this cluster collapsed onto the earliest germ cell clusters rather than grouping with the late-stage cells from which it was isolated (fig. S6A, B), indicating that its transcriptome is nearly identical to that of a pole cell despite its advanced developmental age. Indeed, a previous single-cell atlas of *Drosophila* embryos also observed a population of late-stage germ cells retaining a maternal transcriptome (*25*). Our data reveal that these germ cells fail specification.

We can identify these transcriptionally stalled cells *in vivo*. HCR *in situ* hybridization coupled with immunofluorescence revealed a population of germ cells in wild-type embryos that retained maternal transcripts and failed to activate transcription (Fig. 5A, B). During gastrulation, all germ cells appeared homogeneous, each containing detectable *nanos* RNA (Fig. 5A). At the beginning of migration, however, the germ cell cluster resolved into two transcriptionally distinct populations: those that retained high levels of maternal transcripts such as *nanos*, and those that began degrading their maternal transcriptome. These populations were also spatially distinct, with the maternal-retaining cells remaining in the gut while their counterparts migrated toward the gonad. Notably, the maternal-transcript-retaining cells showed reduced Vasa protein compared to their migrating counterparts, suggesting that translation of maternal mRNAs may also be compromised in this population (Fig. 5A). By stage 11, these two populations were both transcriptionally distinct and spatially separated within the embryo. *in situ* hybridization for *Taf12L*, which marks transcriptionally active germ cells, revealed that the gut-retained population had no detectable zygotic transcription, while healthy migrating germ cells robustly expressed *Taf12L* (Fig. 5B). Rather than migrating to the gonad with their healthy counterparts, the transcriptionally stalled cells accumulated within the gut where they ultimately died (Fig. 5C). Because these germ cells were lost during their migration to the gonad, we call them the lost children. At peak abundance, an average of 10 lost children were present per embryo, but these cells progressively died within the gut, and by late gonad coalescence no detectable germ cell signal remained (Fig. 5D).

**Figure 5:**
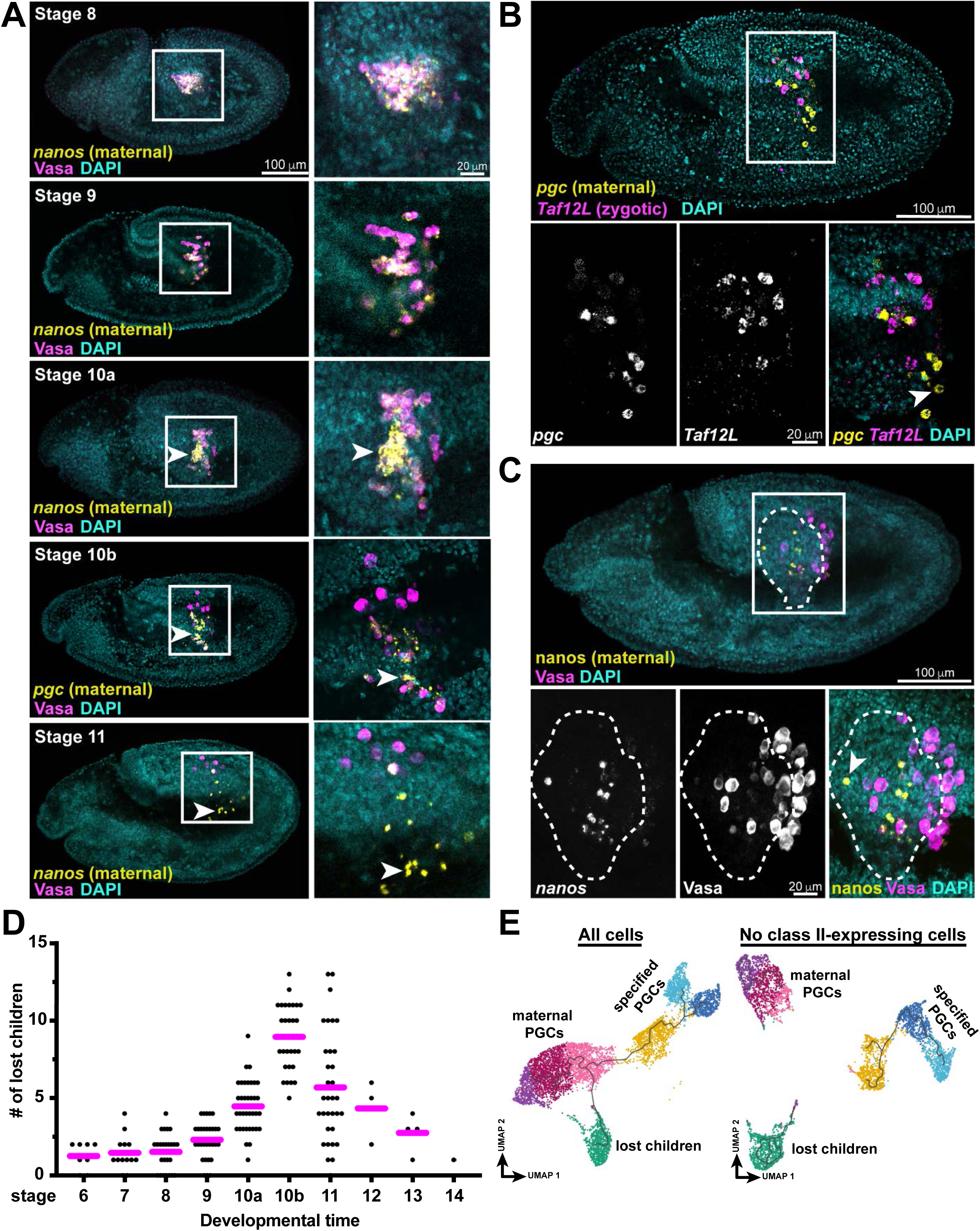
A subset of primordial germ cells fails to activate the totipotency program and dies. A. HCR *in situ* hybridization combined with immunofluorescence of germ cells from wild-type embryos throughout development, showing Vasa protein (magenta) and the maternal transcripts *nanos* or *pgc* (yellow). Full embryo (left) with germ cell inset (right). Arrowheads mark lost children, which fail to degrade maternal transcripts. Nuclei marked with DAPI (cyan). Scale bars, 100 µm (full embryo) and 20 µm (inset). B. HCR *in situ* hybridization of a stage 10 wild-type embryo (top) with high-magnification imaging of germ cells (bottom). *Taf12L* (magenta) is present in healthy germ cells and absent from lost children. *pgc* (yellow) is maternally deposited and retained in lost children. Arrowheads mark lost children. Nuclei shown with DAPI (cyan). Scale bars, 100 µm (full embryo) and 20 µm (inset). C. Lost children fail to migrate. HCR *in situ* hybridization of a stage 11 wild-type embryo (top) with high-magnification imaging of germ cells (bottom). *nanos* (yellow) marks lost children, which are retained within the gut, shown within the dashed line based on endoderm marker expression. Vasa protein (magenta) marks healthy germ cells that have successfully migrated out of the gut. Nuclei marked with DAPI (cyan). Arrowheads mark lost children. Scale bars, 100 µm and 20 µm. D. Quantification of the number of lost children per wild-type embryo across developmental stages. Each dot represents one embryo, n = 188. E. Bioinformatic removal of cells expressing appreciable levels of Class II transcripts breaks the pseudotime trajectory, separating the maternal pole cells from the specified PGCs.

Across species, a substantial portion of germ cells are lost between formation and gonad coalescence. This is a conserved feature of germline development, occurring in *Drosophila*, Xenopus, and mouse (*11, 13–15, 50*). Despite being observed for decades, the identity and fate of these lost cells had remained unknown. We find that lost children represent 30-40% of total PGCs at late embryonic stages, both *in vivo* and in our scRNA-seq dataset, a proportion consistent with this conserved attrition. These data suggest that lost children are the long-observed but uncharacterized population of germ cells that fail to reach the gonad. This proportion is not determined solely by germ plasm inheritance, as *oskar* NLS mutants, which form fewer pole cells while maintaining the same total germ plasm, still produce lost children (fig. S6C, D) (*15, 51*).

Because lost children ultimately die, we asked whether their death leads to their failed transcription. We examined lost children in *chk2* mutants, which lack p53-mediated apoptosis. In *chk2* mutants, lost children survived but still failed to migrate or degrade maternal transcripts (fig. S6E, F). This shows that their death is a consequence of failed specification, not its cause.

The failure to activate zygotic transcription, detectable at the onset of germ cell specification, is a defining feature of lost children. Within the newly formed germ cell cluster, approximately 40% of pole cells failed to significantly activate Class II gene expression, as quantified by *Taf12L* staining *in vivo* and confirmed across Class II genes in our scRNA-seq dataset (fig. S6G). This proportion matched the fraction of germ cells that ultimately became lost children, suggesting that the distinction between healthy PGCs and lost children is established at the earliest moment of specification. Consistent with this, removing Class II-expressing cells from our scRNA-seq dataset and repeating pseudotime analysis fragmented the single healthy germ cell trajectory into three disconnected populations: newly formed pole cells, healthy specified PGCs, and lost children (Fig. 5E). This indicates that Class II-expressing cells form the bridge connecting maternal pole cells to fully specified PGCs, and that without them the developmental continuum between pole cell and PGC is broken.

Because Class II genes constitute the founding transcriptional program of the totipotent blastomeres, their activation in nascent germ cells marks the establishment of a totipotent chromatin state. This distinction defines two fates among the cells that form at the posterior pole. Pole cells that activate the totipotency program become competent primordial germ cells, and those that fail to activate it are ultimately lost. Establishing this totipotent state, rather than producing functional protein from it, is therefore the defining event of germ cell specification, distinguishing a true primordial germ cell from a pole cell.

## Discussion

Our findings reveal that germ cell identity is established through the activation and stabilization of a totipotent chromatin state. Newly formed germ cells remain largely transcriptionally repressed, consistent with the prevailing model in which global RNA polymerase II repression preserves germline identity (*8, 52*). Yet, some genes escape this repression. We find that nascent germ cells activate a transcriptional program immediately after formation; the same founding program that defines the totipotent blastomeres of the early embryo. We propose that the critical feature of this program is not the transcription itself, which is limited to roughly thirty genes, but the underlying open, permissive chromatin state that germ cells adopt and maintain. Transcription of the Class II genes may be a consequence of this chromatin state rather than its purpose. Because this permissive state allows the transcription of Class II genes, most of which are somatic determinants, germ cells are left poised to differentiate inappropriately. To protect against this, germ cell identity must be secured at the translational level, where Nanos prevents the transcribed program from determining cell fate. Germline identity is therefore established not by what germ cells transcribe, but by the chromatin state they maintain and the translational control that holds its consequences in check, decoupling transcription from cell fate commitment.

This model resolves a long-standing puzzle in germ cell biology: why are the conserved determinants of germline identity RNA-binding proteins rather than transcription factors? Somatic lineages are specified by master-regulator transcription factors that restrict chromatin and drive lineage-specific programs. Germ cells, however, do the opposite. Their identity depends on maintaining an unrestricted, permissive chromatin state antithetical to the ordered chromatin of a differentiated cell. A transcriptional regulator would impose a specific transcriptional program, the very outcome germ cells must avoid. Their fate must therefore be secured post-transcriptionally, leaving the chromatin itself uncommitted. Germ cells accomplish this using RNA-binding proteins such as Nanos, which stabilize the totipotent state by controlling which transcripts are translated. We propose that RNA-binding proteins are the master regulators of germ cell fate across organisms, stabilizing the inherently unstable totipotent chromatin state of the germline throughout development. This explains why the loss of germline RNA-binding proteins, rather than transcription factors, causes germ cells to lose their identity. This framework also explains why in the absence of Nanos, *Drosophila* germ cells differentiate into somatic cell types, adopting the fate of the somatic lineage they lie closest to (*46, 53*). Because germ cells maintain the uncommitted, permissive chromatin of a totipotent blastomere, they are inherently poised to differentiate. In the absence of Nanos, germ cells behave as the totipotent cells they resemble, differentiating according to their location in the embryo (*53*).

In the syncytial blastoderm, Zelda has a dual role. As a pioneer factor, Zelda opens the permissive chromatin state of the totipotent blastomeres, but also drives the somatic differentiation programs of the early embryo (*33, 35*). Zelda dosage is critical, as both too little and excessive Zelda activity cause embryonic lethality (*54*). Germ cells appear to retain sufficient levels for Zelda’s chromatin-opening activity to establish a totipotent blastomere-like state, while using Nanos to prevent Zelda from driving somatic programs forward. This dual logic is conserved in the mammalian germline. TFAP2C and PRDM14 both help maintain the open, pluripotent chromatin state of the naive epiblast and embryonic stem cells, and these two factors also regulate primordial germ cell fate (*55–59*). Together, these observations suggest that germ cells across animals are built around a shared totipotent chromatin state.

Our findings suggest a model for how and when germ cells first establish this totipotent chromatin. In *Drosophila*, as nuclei migrate to the posterior pole and enter the germ plasm, they carry with them the chromatin-opening activity of Zelda that is already active in the surrounding syncytium. If the totipotent chromatin state is established before the nuclei migrate into the germ plasm, then the failure of the lost children may originate there. It would be informative to determine whether this failure is specific to the germline, or whether some soma-fated blastomeres similarly fail to establish the totipotent state. This route may distinguish preformation species such as *Drosophila* from mammals. In the fly, germ cells inherit the open chromatin of the totipotent blastomeres directly, and need only to maintain it. In mammals, primordial germ cells instead arise by induction from epiblast cells that have already begun to differentiate and organize their chromatin. It has long been known that these cells reactivate the core pluripotency network, including Sox2 and Nanog alongside continued Oct4 expression (*60*), but why they do so has been unclear. Our findings suggest that this reactivation re-establishes the same open, totipotent chromatin state that preformation germlines inherit intact. In both cases, the endpoint is a germline that maintains the totipotent chromatin of the early embryo, whether preserved by inheritance or rebuilt by re-establishment. In some species, germ cells are specified at a later stage in development and germ cells can be derived from somatic cells in mammals (*61*). Notably, the earliest germline-specific markers in this transition are translational regulators like Nanos (*59, 60*). We propose that general reprogramming, coupled with Nanos-induced stabilization, enables germ cell specification at any developmental stage.

Our findings also shed new light on a long-standing and unexplained observation in germ cell biology. Across animals, a substantial fraction of primordial germ cells is lost between their formation and gonad coalescence, yet the reason for this loss has remained unknown (*11–15*). We find that this attrition has a transcriptional origin. The lost germ cells fail to activate the totipotency program at the earliest step of the germline maternal-to-zygotic transition. This failure defines a discrete molecular checkpoint. Pole cells that activate the Class II program become competent primordial germ cells that reach the gonad, whereas those that fail to activate it remain in a maternal state, fail to migrate, and are eliminated. The distinction between a pole cell and a specified germ cell is therefore not merely positional or morphological, but transcriptional. These cells also fail to fully clear their maternal transcriptome, a process that is coupled to transcriptional activation during the maternal-to-zygotic transition in somatic cells (*10*). Whether failure to clear maternal transcripts causes or follows failure to activate transcription remains an open question.

Together, our findings redefine how germ cell identity is established. Rather than being specified by a dedicated transcriptional program, germ cells maintain the uncommitted, permissive chromatin of the totipotent blastomeres and transcribe their founding program, but secure their identity post-transcriptionally, through the RNA-binding protein Nanos. This strategy allows germ cells to hold two seemingly incompatible states at once: the differentiated identity of a specified lineage and the totipotent potential of the early embryo. In this way, germ cells decouple what they transcribe from what they become. We propose that this decoupling, rather than any single germline determinant, may provide the conserved foundation of germline identity.

## Materials and Methods

### Fly Stocks and Husbandry

All stocks and crosses were maintained at 18 °C and 25 °C, on cornmeal molasses yeast medium. The *Taf12L* knockout and *Amos-GFP* lines were generated for this study as described below. Lehmann lab *w1118*, *P[vas-GFP], nos^L7^, nos^BN^, osk-nls* mut stocks were used throughout the study. The *zelda shRNA* stock was a gift from C. Rushlow. The Chk2 mutant stock (*loki^p6^*) was a gift from Paul Lasko. For all *nanos* mutant experiments, *nos^BN^* and *nos^L7^* flies were crossed to yield F1 offspring that were either heterozygous for the *nos^BN^* or *nos^L7^* allele or transheterozgous *nos^BN^*/*nos^L7^* mutants. F1 control heterozygous and *nos^BN^*/*nos^L7^* mutant females were mated to produce F2 *nos* maternal heterozygous and *nos* maternal mutant embryos that were used for downstream experiments. *W1118* was used as a wild-type control. For Zelda knockdown experiments, females carrying the oogenesis-specific driver, *mata-Gal4-VP16* were crossed to *UAS-Zelda-shRNA* males to yield F1 offspring with one copy of the driver and one copy of the *zelda*-shRNA. This ensured expression of the *zelda shRNA* in mid-late oogenesis (stages 5-14) in F1 females and strong reduction of maternal *zelda* in F2 embryos. *mata-Gal4-VP16* embryos were used as controls. For Chk2 experiments, homozygous *loki^p6^* mutant females were crossed to homozygous *loki^p6^* mutant males (as these flies are viable and fertile), to generate *loki^p6^* maternal and zygotic mutant embryos.

### Embryo collection timing for transcriptomic atlas

Embryos were collected from four overlapping developmental windows, each defined by a collection period followed by an aging step. The first timepoint used a 1-hour collection and a 1-hour aging step, so that processing began when embryos were 1-2 hours post fertilization (hpf). The second collection used a 2-hour collection window and a 1.5-hour aging step (1.5-3.5 hpf at the start of processing). The third used a 3-hour collection window and a 3.5-hour aging step (3.5-6.5 hpf at the start of processing), and the fourth used a 3-hour collection and a 6.5-hour aging step (6.5-9.5 hpf at the start of processing). Images in fig. S1A show a sample of embryos at the start of processing. Because subsequent dissociation and FACS took a further 1.5 to 3 hours, during which time development can continue, each timepoint spans a range of developmental stages from the start of processing through the end of sorting (1.5-4, 1.5-6, 4-9, and 7-12 hpf). Analysis revealed that the germ cells from the first two timepoints occupied a completely overlapping range of pseudotime. We therefore treated these two timepoints as a single specification timepoint for the second replicate.

The early timepoints contained a small proportion (approximately 5-10%) of older embryos, consistent with occasional egg retention by *Drosophila* females. Germ cells from these embryos were readily identified during analysis, as they clustered with the later timepoints rather than with the cells from their collection window, and were retained at their correct developmental position within the atlas. The lost children are distinct from this collection contamination. The youngest embryos possible in the latest collection window were 6.5 hours old (approximately stage 10-11), yet the lost children isolated from this window retain a fully maternal transcriptome resembling that of 2-4 hour-old embryos. This transcriptome is far younger than any embryo that could have been present in the collection, suggesting that the lost children reflect a genuine developmental failure rather than contamination by younger embryos. In our collections, we also did not observe embryos that had arrested early in development, ruling out developmental arrest as a source of the lost children.

### Embryo Single cell suspension and germ cell isolation by FACS

Embryos were collected from population cages maintained in a 25°C incubator, aged for the appropriate length of time in the incubator, and dechorionated in 50% bleach for 5 min on a nutator (60 rpm), then rinsed thoroughly with water in a collection basket. Dechorionated embryos were transferred into ice-cold FACS buffer (55 mM NaCl, 40 mM KCl, 5 mM CaCl2, 15 mM MgSO4, 10 mM Tricine pH 7.0, 20 mM glucose, 50 mM sucrose, and 1 g/L BSA, final pH 7.0) and dounce-homogenized on ice in batches (approximately 20-25 strokes) until no visible chunks remained. Homogenized samples were centrifuged at 800 × g for 3 min at 4°C, and the pellet was washed with FACS buffer and re-centrifuged until the supernatant ran clear (typically two washes). All steps were performed on ice with pre-chilled buffer, and wide-bore (P1000) tips were used throughout to minimize shear.

Dead cells were depleted using the Akadeum Dead Cell Removal Microbubble Kit (Akadeum Life Sciences), according to manufacturer’s protocol. To label live cells, the final cell pellet was resuspended in 1-2 mL of FACS buffer and cell concentration was estimated using a Countess automated cell counter (Thermo Fisher). Cells were adjusted to approximately 5 × 10^6 cells/mL, and Calcein Violet AM (Invitrogen, cat. no. C34858) was added according to the manufacturer’s instructions. Labeled cells were filtered into FACS tubes and kept on ice until sorting.

Live germ cells were isolated on a FACS Aria (BD Biosciences) at the Whitehead Institute Flow Cytometry Core. Cells were gated sequentially on forward and side scatter to exclude debris, on FSC-H versus FSC-A and SSC-W versus SSC-A to select single cells, on Calcein Violet fluorescence to select live cells, and on GFP fluorescence to isolate the P{vasa-GFP}-positive germ cell population (fig. S1B). Sorted germ cells were collected into Schneider’s Drosophila medium (Gibco, cat. no. 21720024) supplemented with 10% FBS for downstream single-cell RNA-sequencing.

### Single-cell RNA-sequencing and data processing

Live vasa:GFP-positive germ cells were isolated by FACS and processed for single-cell RNA-sequencing using the 10x Genomics Chromium Single Cell 3’ kit (v3). Libraries were sequenced on an Illumina NovaSeq using an S4 flow cell (150 × 150 bp paired-end reads). Reads were mapped to the dm6 reference genome built to include both mRNA and some lncRNA annotations. Reads were mapped using the 10X CellRanger pipeline with default parameters, on a per-sample basis. A total of 15,712 cells were captured across the four timepoints. Downstream analysis was performed in R using Seurat. Each timepoint was first processed independently for quality control. Cells with more than 200 detected genes and fewer than 5,000-6,000 detected genes were retained for analysis. Principal component analysis was performed and cells were clustered using a shared-nearest-neighbor graph and visualized by UMAP. Contaminating somatic clusters were identified and removed based on somatic marker expression (including *elav*, *nrv2*, *slam*, and *nullo*), and dying cells by apoptotic markers (*hid* and *Corp*). The remaining germ cells were re-processed. The four quality-controlled timepoints were then merged into a single dataset and re-processed as above. The final germ cell atlas contained 7,355 cells spanning embryogenesis.

For pseudotime analysis, the processed Seurat object was imported into Monocle3 using SeuratWrappers (incorporating the bug fix from commit 02754e1; github.com/tfrayner). Cells were clustered (cluster_cells, resolution 1e-5), the principal graph was learned (learn_graph), and cells were ordered (order_cells) with the root selected as the node centered in cells expressing a maternal deposition signature, defining the pseudotime trajectory. To determine genes that vary over pseudotime, the Monocle3 graph_test function was used with a modification: the called subfunction calculateLW was edited to replace Matrix::rBind in line 93 to rbind. Genes that vary significantly over pseudotime were defined by a q value of 0 and Moran’s I > 0.25. Counts over pseudotime for each gene were extracted using the Monocle3 exprs function, smoothed using the base R smooth.spline function, and Z-normalized by dividing the mean of each gene by its standard deviation. The heatmap was produced using ComplexHeatmap and the pdf was generated with the aid of the R package magick.

### Expression dynamics of transcriptional classes across pseudotime

Genes were assigned to one of four transcriptional classes (Class I, maternal; Class II, early zygotic; Class III, minor ZGA; Class IV, major ZGA) based on genes that vary over pseudotime as described above. Because some Class II genes are expressed transiently and stochastically, their low overall variance placed them below the threshold for computational detection. These were manually curated by identifying transcripts with expression profiles matching those of known Class II genes, such as *amos*. To visualize the expression dynamics of each class across developmental time, pseudotime values were extracted from the Monocle3 cell dataset object using the pseudotime() function. For each cell, the mean normalized expression of all genes within a given class was calculated from the log-normalized expression matrix. Per-class mean expression was then scaled to a range of 0 to 1 across all cells to allow comparison of relative dynamics between classes of differing absolute expression levels. Scaled expression was plotted against pseudotime for each class, and trends were visualized using locally estimated scatterplot smoothing (LOESS; span = 0.3) in ggplot2. The four classes displayed sequential dynamics consistent with a maternal-to-zygotic progression, with Class II genes showing a transient peak during the specification window.

### Trajectory analysis following removal of Class II-expressing cells

To assess the contribution of Class II-expressing cells to the continuity of the germ cell developmental trajectory, cells with high Class II expression were filtered from the earliest timepoints and the trajectory was recomputed. Within the two earliest timepoints, cells were classified as Class II-expressing and removed if their summed normalized Class II expression exceeded 5 and their *Taf12L* expression exceeded 0.57, while the maternal and lost children populations were preserved. The remaining cells were re-processed through the Monocle3 trajectory-inference pipeline (preprocess_cds, reduce_dimension, cluster_cells, and learn_graph), and the resulting trajectory was compared to that of the full atlas.

### Harmony batch correction of the germ cell atlas

To assess the transcriptional similarity of the lost children to newly formed germ cells, batch correction was performed on the germ cell atlas using Harmony (*62*), which integrates cells across batches and timepoints by iteratively adjusting their embeddings to remove technical variation while preserving biological structure. Following Harmony integration, the lost children clustered with the earliest germ cells rather than with the late timepoint from which they were isolated.

### Embryo collection and fixation for imaging

Flies were kept overnight in cages on apple juice agar plates smeared with yeast paste. Plates were collected and yeast paste removed, and embryos were dechorionated in 50% bleach for 5 min with rocking. Dechorionated embryos were washed onto a mesh within a collection basket, rinsed thoroughly with water, and transferred into a scintillation vial containing 5 mL of 4% paraformaldehyde in PBS and 5 mL heptane. Samples were fixed on a rocker for 20 min, after which the lower aqueous paraformaldehyde layer was removed completely with a glass Pasteur pipette. Methanol (10 mL) was added and the vial was shaken for 30 s to crack and remove the vitelline membranes. Devitellinized embryos sank to the bottom of the vial and were collected with a cut pipette tip. Embryos were washed three times in methanol and stored at −20°C for later use.

### RNA in situ hybridization by hybridization chain reaction (HCR)

RNA in situ hybridization was performed by hybridization chain reaction using either HCR^TM^ v3.0 or HCR^TM^ HiFi Gold (Molecular Instruments), according to manufacturer’s protocols. All probes, buffers, and amplifiers were obtained from Molecular Instruments. For both 3.0 and Gold protocols, methanol-stored embryos were rehydrated through a graded methanol/PBST series (25%, 50%, 75%, and 100% 0.3% Triton X-100 in PBS) and permeabilized in 2.5% Triton X-100 in PBS overnight at 4°C on an orbital rocker.

For HCR v3.0, permeabilized embryos were rinsed twice in 0.3% PBST and washed twice for 10 min in 5× SSCT. Embryos were post-fixed in 4% paraformaldehyde in PBS for 10 min, rinsed three times in PBST, and washed twice for 10 min in 5× SSCT. Samples were pre-hybridized in 200 µL of hybridization buffer (Molecular Instruments) at 37°C until equilibrated. Hybridization buffer was then replaced with 125 µL of fresh buffer containing probe. Probe concentration was determined empirically according to target transcript abundance: on average, 1-3 µL of 1 µM probe was used per sample, with 3 µL required for consistent detection of early zygotic transcripts and 1 µL sufficient for abundant germline markers such as *nanos* and *pgc*. Samples were incubated for two days at 37-42°C in a thermomixer (500 × g). Excess probe was removed by washing four times for 15 min at 37°C in probe wash buffer (Molecular Instruments), followed by a 10 min wash in 5× SSCT. Samples were equilibrated in 200 µL of amplification buffer (Molecular Instruments) at room temperature. Hairpin amplifiers were snap-cooled (95°C for 90 s, then cooled in the dark to room temperature for 30-120 min), and amplification buffer was replaced with 125 µL of fresh buffer containing 1.25 µL of each hairpin. Amplification proceeded overnight at 25°C on an orbital rocker in the dark. Samples were then washed in 5× SSCT at room temperature in the dark for one day, exchanging buffer every 2 h, followed occasionally by an overnight wash in 5× SSCT containing 2 µg/mL DAPI at 4°C for samples with high background. Samples were washed a final 2 h in 5× SSCT and equilibrated in mounting medium (SlowFade Gold, Invitrogen) overnight before mounting.

For HCR HiFi Gold, permeabilized samples were pre-hybridized in HCR HiFi Probe Hybridization Buffer for 30 min at 37°C, then hybridized with probes (1-4 µL of each probe per 200 µL hybridization buffer) for at least 3 h (or overnight) at 37°C. Excess probe was removed by washing four times for 15 min in HCR HiFi Probe Wash Buffer at 37°C. Samples were pre-amplified in HCR Gold Amplifier Buffer for 30 min at room temperature. Hairpins h1 and h2 were snap-cooled separately (95°C for 90 s, then cooled to room temperature in the dark for 30 min), combined into HCR Gold Amplifier Buffer, and applied to the samples. Amplification proceeded for at least 3 h (or overnight) at room temperature in the dark on a nutator. Excess hairpins were removed by washing four times for 15 min in HCR Gold Amplifier Wash Buffer at room temperature, with DAPI added to the second to last wash. Samples were stored at 4°C protected from light and mounted in SlowFade Gold Antifade Mountant with DAPI (Invitrogen, cat. no. S36936) prior to imaging. All samples were imaged on a Zeiss LSM 980 confocal microscope.

### Combined immunofluorescence and RNA in situ hybridization

For simultaneous protein and RNA detection, immunofluorescence was combined with HCR RNA in situ hybridization, according to manufacturer’s protocols (Molecular Instruments). All antibody encoders were purchased from Molecular Instruments. Briefly, Methanol-stored embryos were rehydrated through a graded methanol/PBST series (75%, 50%, 25% methanol in 0.3% Triton X-100 in PBS, 5 min each), washed in 0.3% PBST, and permeabilized in 2.5% Triton X-100 in PBS overnight at 4°C. For immunofluorescence, embryos were blocked in HCR HiFi Antibody Buffer (Molecular Instruments) for 4 h at 4°C, then incubated with primary antibody diluted in antibody buffer overnight at 4°C (antibodies and dilutions listed in the key resources table). Samples were rinsed and washed 4 × 30 min in 0.3% PBST, then incubated overnight at 4°C with HCR-hairpin-conjugated encoders (Molecular Instruments) matched to the host species of the primary. The hairpin on the encoder was subsequently detected by HCR amplification. Samples were then rinsed and washed 3 × 10 min in 0.3% PBST, washed in 5× SSCT (0.1% Tween-20), and post-fixed in 4% paraformaldehyde for 10 min at room temperature, then rinsed 3 × 10 min in 0.3% PBST and washed 2 × 10 min in 5× SSCT. RNA detection was then performed by HCR exactly as above.

### Immunofluorescence

Methanol-stored embryos were rehydrated through sequential 5 min washes of 25%, 50%, and 75% methanol in 0.3% Triton X-100 in PBS (PBST) and blocked in blocking buffer (PBST with 1% BSA) for 30 min. Primary antibodies were diluted in blocking buffer and incubated overnight at 4°C (antibodies used in this study are listed in the key resources table). Samples were washed 4 × 15 min in PBST, blocked again for 30 min, and incubated with secondary antibodies (1:500 in blocking buffer) overnight at 4°C with rocking. Samples were washed 4 × 15 min in PBST, with DAPI added to the penultimate wash, and equilibrated in mounting medium (SlowFade Gold, Invitrogen) overnight before mounting. Samples were imaged on a LSM 980 confocal microscope.

In some experiments, conventional immunofluorescence was performed prior to HCR RNA *in situ* hybridization. Embryos were rehydrated and permeabilized as described, blocked, and incubated with primary antibody overnight at 4°C, followed by a standard fluorophore-conjugated secondary antibody. Following antibody staining, samples were post-fixed in 4% paraformaldehyde for 10 min at room temperature to preserve the immunofluorescence signal, and HCR RNA *in situ* hybridization (v3.0 or Gold) was then performed as described above.

### Posterior-pole mounting of embryos

To visualize primordial germ cells at the posterior pole, stained embryos were transversally bisected and mounted upright. Embryos in antifade mounting medium were pipetted onto a slide using a cut pipette tip, and excess mounting medium was wicked away with a Kimwipe. Under a stereomicroscope, embryos of the desired stage were aligned in a single row with their posterior poles oriented in one direction. Using a scalpel, the posterior third of each embryo was excised with a single vertical cut, and the anterior portions were discarded. The posterior fragments were then oriented cut-side down to position the germ-cell-containing posterior pole facing the objective. Excess moisture was removed as needed to help the fragments remain upright. A coverslip was gently lowered onto the aligned fragments, additional mounting medium was added at the coverslip edges, and the coverslip was sealed at the corners with clear nail polish and allowed to dry before imaging.

### Chromatin accessibility analysis of *Drosophila* snATAC-seq data

Pseudobulk single-nucleus ATAC-seq (sci-ATAC-seq3) bigWig tracks for the Drosophila melanogaster embryo were obtained from Calderon et al. (GEO: GSE190130), which were split by developmental time windows and cell clusters (*41*). Cell type annotations for each track were assigned based on the supplementary information of the original paper. Tracks annotated as “unknown” were excluded from downstream analysis.

To study the genome-wide chromatin accessibility similarity between cell types, the Drosophila genome (version dm6) was binned into 5 kb intervals, and the mean ATAC signal in each bin was extracted from every bigwig track with pyBigWig. Pairwise similarity between tracks was measured as the Pearson correlation of these per-bin mean signal vectors. Similarity matrices were computed independently for each two-hour time window (12-14 hours for late stage germ cells and 2-4 hours for newly formed germ cells), and restricted to the cell types present in that window. We averaged blastoderm clusters from the earliest windows (0-2 hours), to form a single “totipotent blastomeres” vector, which was included in every per-window matrix so that all windows were compared against a common blastoderm reference. Correlation matrices were converted to distance matrices (1 − correlation), and hierarchical clustering was performed with average linkage. Heatmaps were generated in R using ComplexHeatmap, with cells colored by correlation coefficient on a continuous scale from 0.5 to 1.

To further characterize the enrichment of open chromatin around promoter regions or peak regions, two metrics including TSS enrichment and Fraction of Signal in Peaks (FSiP), were used for quantification. TSS enrichment was computed following the ENCODE standard (*63*) (https://www.encodeproject.org/data-standards/terms/#enrichment). Fraction of signal in peaks was defined as the total signal within consensus ATAC peaks divided by the total signal on autosomes. The consensus peak set was downloaded from the original paper. For both metrics, the sex chromosomes were excluded to avoid confounding by dosage compensation, and the mitochondrial DNA was also excluded to remove contamination. TSS enrichment and FSiP for each cell type were obtained by averaging across all the tracks of this cell type.

### Generation of the amos-GFP line

The *amos-GFP* line was generated by CRISPR/Cas9-mediated homology-directed repair. A single guide RNA (CTTCAGATTTCCACTAGGATATGA) was cloned into the pU6.3 gRNA vector. The repair template was based on the pHD-ScarlessDsRed donor vector (DGRC #1364). A 1,015 bp homology arm upstream of the *amos* +1 site was cloned upstream of the 3xP3-dsRED marker cassette, and a 1,000 bp downstream homology arm beginning at the *amos* stop codon was cloned downstream of the marker. A linker and superfolder GFP (sfGFP) tag were inserted in frame between the end of the *amos* coding sequence and the stop codon, and the *amos* 3’UTR was cloned downstream of the GFP tag and upstream of the 3xP3-dsRED cassette. To facilitate cloning, a segment spanning the last 19 bp of the sfGFP sequence and the first 50 bp of the *amos* 3’UTR was inserted together with the 3’UTR. A point mutation was introduced to alter the PAM site from ACC to AGC to prevent re-cutting of the repaired locus. Guide and repair plasmids were injected by BestGene into embryos expressing vas-Cas9 (Bloomington Drosophila Stock Center #51324). Four edited founder males were isolated, the vas-Cas9 transgene was removed by outcrossing, and balanced stocks were established. The 3xP3-dsRED cassette was retained, allowing the edited allele to be tracked by dsRED expression.

### Generation of the Taf12L knockout line

Two independent *Taf12L* knockout lines were generated using CRISPR/Cas9-mediated homology-directed repair, following an approach similar to that previously described (Grill et al., 2023) (*19*). The two lines were made using identical repair plasmids but distinct sets of guide RNAs targeting different sites within the locus, providing an independent control against off-target effects. Guide RNAs were identified using the flyCRISPR Target Finder (http://targetfinder.flycrispr.neuro.brown.edu/), and for each line two gRNAs (sequences listed in the key resources table) were cloned into the pCFD4 plasmid. A pScarless repair plasmid was constructed with homology arms comprising approximately 1 kb immediately upstream of the *Taf12L* 5’UTR and approximately 1 kb immediately downstream of the *Taf12L* 3’UTR, such that the entire *Taf12L* locus (from the 5’UTR through the 3’UTR) was replaced by a dsRED cassette. The dsRED cassette was not excised, allowing mutant animals to be tracked by dsRED expression. Plasmids were injected by BestGene into embryos expressing vasa-Cas9 (Bloomington Drosophila Stock Center #51324) using standard procedures.

### Quantitation of Zelda, RNAPII S5, and S2 signal

Nuclear immunofluorescence intensity for RNAPIIS2p, RNAPIIS5p, and Zelda was quantified using a custom FIJI/ImageJ macro. For images acquired as z-stacks, a single best-focus z-plane was selected manually to avoid the z-depth artifacts associated with maximum-intensity projection. Nuclei were segmented automatically from the DAPI channel by Gaussian blurring (sigma = 2), adaptive local thresholding (Phansalkar method, radius = 50 px), watershed separation, and particle analysis (minimum size = 30 px^2; nuclei touching image edges excluded); segmentation was reviewed for each image and parameters were adjusted when necessary. Within each embryo, approximately 20 somatic nuclei and the germ cell nuclei were manually classified based on position, morphology, and the presence of a germ cell marker. Mean fluorescence intensity of the target channel (RNAPIIS2p, RNAPIIS5p, or Zelda) was measured within each segmented nuclear region of interest, and the signal in each nucleus was normalized to the mean signal of the somatic nuclei within the same plane in the same embryo to control for staining and imaging variability between embryos. Statistical comparisons were performed at the embryo level in R. Normalized germ cell signal was averaged within each embryo, and mean values were compared between conditions using a Wilcoxon rank-sum test. In scatter plots, each point represents an individual nucleus, while statistical tests were performed on per-embryo means.

### Quantitation of Amos-GFP protein signal in pole cells

To quantify Amos-GFP protein in the germline, confocal z-stacks of control, *nanos* heterozygous, and *nanos* mutant embryos were maximum-intensity projected in FIJI/ImageJ. Display levels in the Amos-GFP channel were adjusted so that true negatives appeared fully black, and the number of Amos-GFP-positive pole cells per embryo was counted manually, with pole cells identified by Vasa staining. Counts were performed independently by two individuals. The number of Amos-GFP-positive pole cells was compared between control and *nanos* mutant embryos using an unpaired Welch’s t-test.

### Quantification of HCR signal across transcriptional waves

HCR in situ hybridization signal was quantified in germ cells using FIJI/ImageJ. Confocal z-stacks were opened in color mode, and germ cells were identified by the P{vasa-GFP} GFP channel. For each germ cell, a polygon was drawn around the cell using only the vasa-GFP channel, and the mean fluorescence intensity of the transcript channel within that region was measured. Background was measured by moving the same polygon to a region of the embryo containing background signal. This was repeated for every germ cell in the embryo. For each cell, transcript signal was background-subtracted, and cells were scored as positive or negative for the transcript based on whether their background-subtracted signal exceeded the threshold. The proportion of transcript-positive germ cells was calculated per embryo.

### Spot detection and quantitation of transcript abundance

Confocal z-stacks acquired as multichannel .czi files were split into single-channel stacks, and the germline marker and DAPI channels were combined into a two-channel stack for segmentation of germ cell boundaries using a locally fine-tuned Cellpose-SAM model (*64–66*). Individual RNA puncta were detected independently in each channel using Spotiflow, a deep-learning-based spot detector that outputs subpixel-accurate coordinates alongside a per-spot detection probability and local intensity (*67*). Detection thresholds for each marker were calibrated on an annotated training dataset using Spotiflow’s built-in optimization routine. Detected spots were mapped to individual germ cells using the segmentation masks, and spots outside labeled cells were excluded. Finally, spot counts per cell were normalized to cell volume to calculate puncta density.

### Quantitation of lost children

The number of lost children per embryo was quantified manually across developmental stages. Confocal images were despeckled in FIJI/ImageJ for noise reduction. Germ cells were identified by Vasa protein staining, and lost children were scored as Vasa-positive germ cells that retained nanos RNA and were positioned closer to, or within, the gut, away from the migrating germ cell population.

### Quantitation of Taf12L expression in pole cells

The proportion of Taf12L-expressing pole cells was quantified manually. Confocal images were despeckled in FIJI/ImageJ for noise reduction. Pole cells were identified by *pgc* RNA staining, and the number of Taf12L-positive pole cells was scored and divided by the total number of pole cells in each embryo to give the percentage of Taf12L-positive pole cells.

## Acknowledgments

We thank R. Young and T. Lee for their intellectual guidance and advice throughout this project; Y. Yamashita for helpful discussions; we thank C. Rushlow for sharing the Zelda antibody and the *zelda shRNA* fly line; P. Lasko for sharing the chk2 mutant fly line; M. Selvaraj for assistance with embryo dissociations and FACS for the scRNA-sequencing experiments; J. Love and the Genome Technology Core at the Whitehead Institute for technical support with scRNA-sequencing experiments; K. Daniels and the Flow Cytometry Core at the Whitehead Institute for technical support; I. Barrasa and the Bioinformatics and Research Computing group (BaRC) at the Whitehead Institute for bioinformatic support; The Whitehead Institute Keck Imaging Facility and specifically A. Marcos Vidal for image analysis, quantitation, and support; and the Lehmann and Yamashita laboratories for helpful discussions. Large language model-based AI assistance (Claude, Anthropic) was used during the preparation of this manuscript for language editing and refinement of text, and debugging code in R and ImageJ. The authors reviewed, edited, and verified all AI-assisted text and code. The authors take full responsibility for the content, analyses, interpretations, and conclusions presented in this work. No AI tool was used to generate, alter, or fabricate data or results.

## Author contributions

S.G. isolated and purified germ cells and performed and analyzed all scRNA-sequencing experiments with input from A.A.R. A.R. and S.G. recognized the lost children phenotype, characterized and quantified lost children and Class II genes *in vivo*, performed HCR *in situ* hybridization and immunofluorescence for the lost children and Class II genes, and quantified the lost children and the proportion of Taf12L-expressing pole cells. C.M. and S.G. recognized that Class II genes are expressed in the blastomeres, and C.M., S.G., and L.V.P. performed HCR *in situ* hybridization of Class II genes in the germline and blastomeres. C.M. generated and validated the Amos-GFP CRISPR line. L.V.P. and S.G. performed HCR *in situ* hybridization and immunofluorescence for Nanos and Class II gene expression and quantified Amos-GFP, Zelda, RNAPIIS5p, and RNAPIIS2p signal. S.G. performed the maternal Zelda knockdown, with conceptual input from A.T. X.G. and S.G. performed the snATAC-seq bioinformatic analysis of data from Calderon et al., with X.G. generating the processed data and S.G. producing the chromatin figures. A.T. stained and imaged embryos for representative transcripts of each class, and A.A.R generated the transcriptional heat map. A.T., C.M., and S.G. quantified the dynamics of each transcriptional class. S.G., A.R., and R.L. conceived the project. R.L. supervised the work. S.G. wrote the manuscript, and all authors discussed the results and commented on the manuscript.

## Data and materials availability

Requests for fly lines generated in this study should be directed to the corresponding author. All sequencing data will be deposited to the Gene Expression Omnibus (GEO) upon publication, and can be sent to reviewers on their request.

---

**fig. S1 (related to Figure 1):**
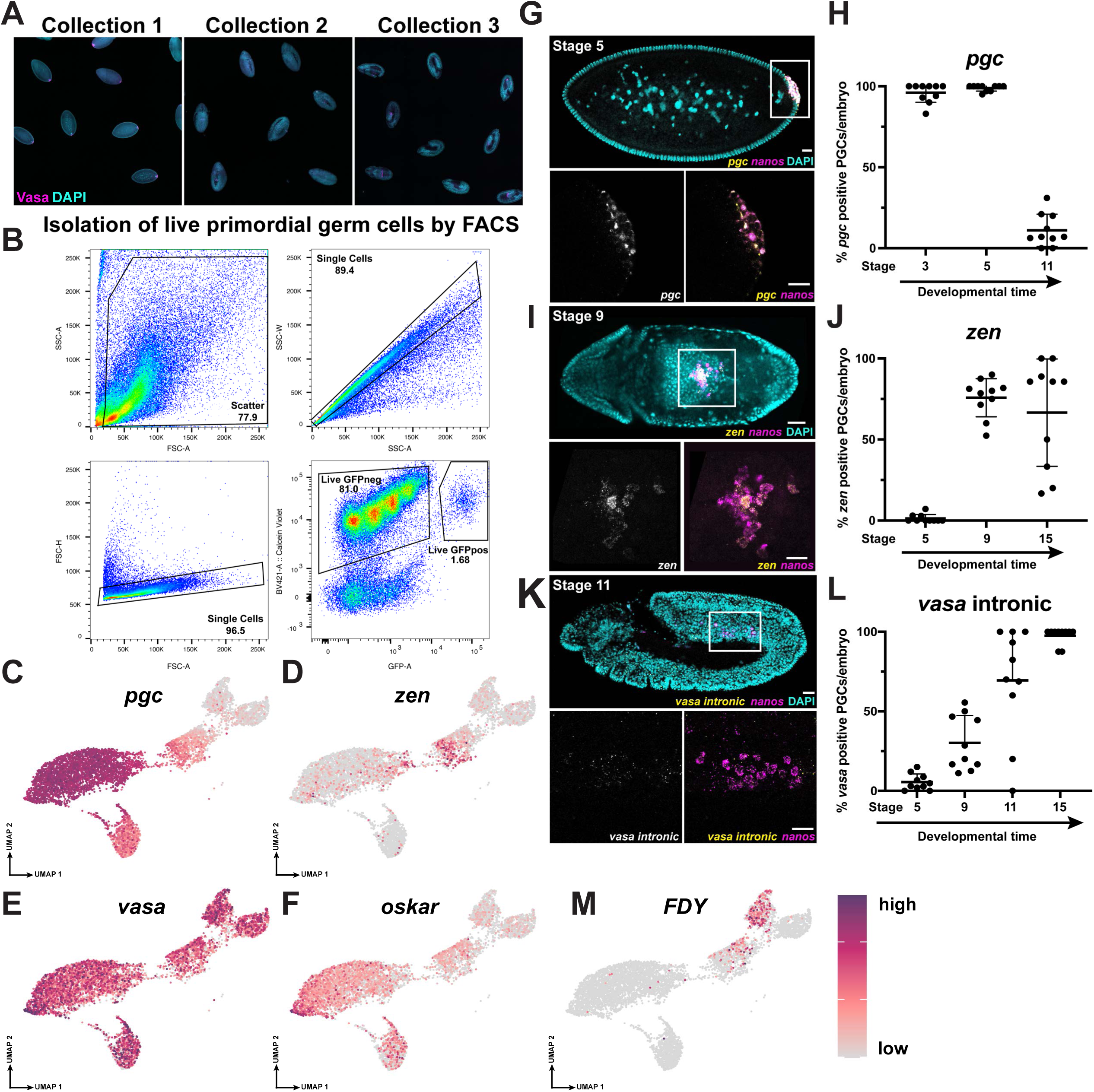
Isolation and validation of the primordial germ cell scRNA-seq atlas. A. A subset of P{*vasa-GFP*} embryos from each of the three collections was stained with DAPI and Vasa to mark the germline and assess the heterogeneity of the collected pool. At the earliest stage, contaminating older embryos constitute approximately 5% of the total, but germ cells from these embryos are readily distinguished bioinformatically (see Fig. 1B). These collections represent embryos prior to dissociation and FACS, approximately 1.5-2 hours before cells were sequenced. B. FACS gating strategy for isolating the live germ cell population from dissociated P{*vasa-GFP*} embryos. Cells were sorted for the presence of GFP and Calcein Violet (live cell marker). C. Feature plot showing expression of the maternal gene *pgc* from scRNA-seq analysis. D. Feature plot showing expression of the zygotic gene *zen* from scRNA-seq analysis. E. Feature plot showing expression of the ubiquitous gene *vasa* from scRNA-seq analysis. F. Feature plot showing expression of *oskar* from scRNA-seq analysis. G. *In vivo* validation of *pgc* dynamics by HCR *in situ* hybridization. *pgc* transcript (yellow), germ cells marked by *nanos* (magenta). Full embryo (top) with high-magnification inset of germ cells (bottom). Nuclei stained with DAPI (cyan). Scale bars, 20 µm. H. Quantification of *pgc*-positive germ cells recapitulates the transcriptional dynamics observed by scRNA-seq. Each dot represents one embryo. I. *In vivo* validation of *zen* dynamics by HCR *in situ* hybridization. *zen* transcript in yellow, germ cells marked by *nanos* transcript in magenta. Nuclei shown with DAPI (cyan). Scale bars, 20 µm. J. Quantification of *zen*-positive germ cells recapitulates the transcriptional dynamics observed by scRNA-seq. Each dot represents one embryo. K. Identification of the onset of zygotic *vasa* transcription by HCR *in situ* hybridization using probes targeting *vasa* intronic sequences. Because *vasa* is both maternally deposited and zygotically transcribed, its zygotic activation cannot be resolved in the scRNA-seq atlas. Intronic probes detect only nascent transcripts. Nascent *vasa* transcript in yellow, germ cells marked by *nanos* transcript in magenta. Nuclei shown in cyan. Scale bars, 20 µm. L. Quantification of the proportion of germ cells with nascent *vasa* transcription across developmental stages. Each dot represents one embryo. M. Feature plot showing expression of the Y-chromosome gene *FDY* from scRNA-seq analysis.

**fig. S2 (related to Figure 2):**
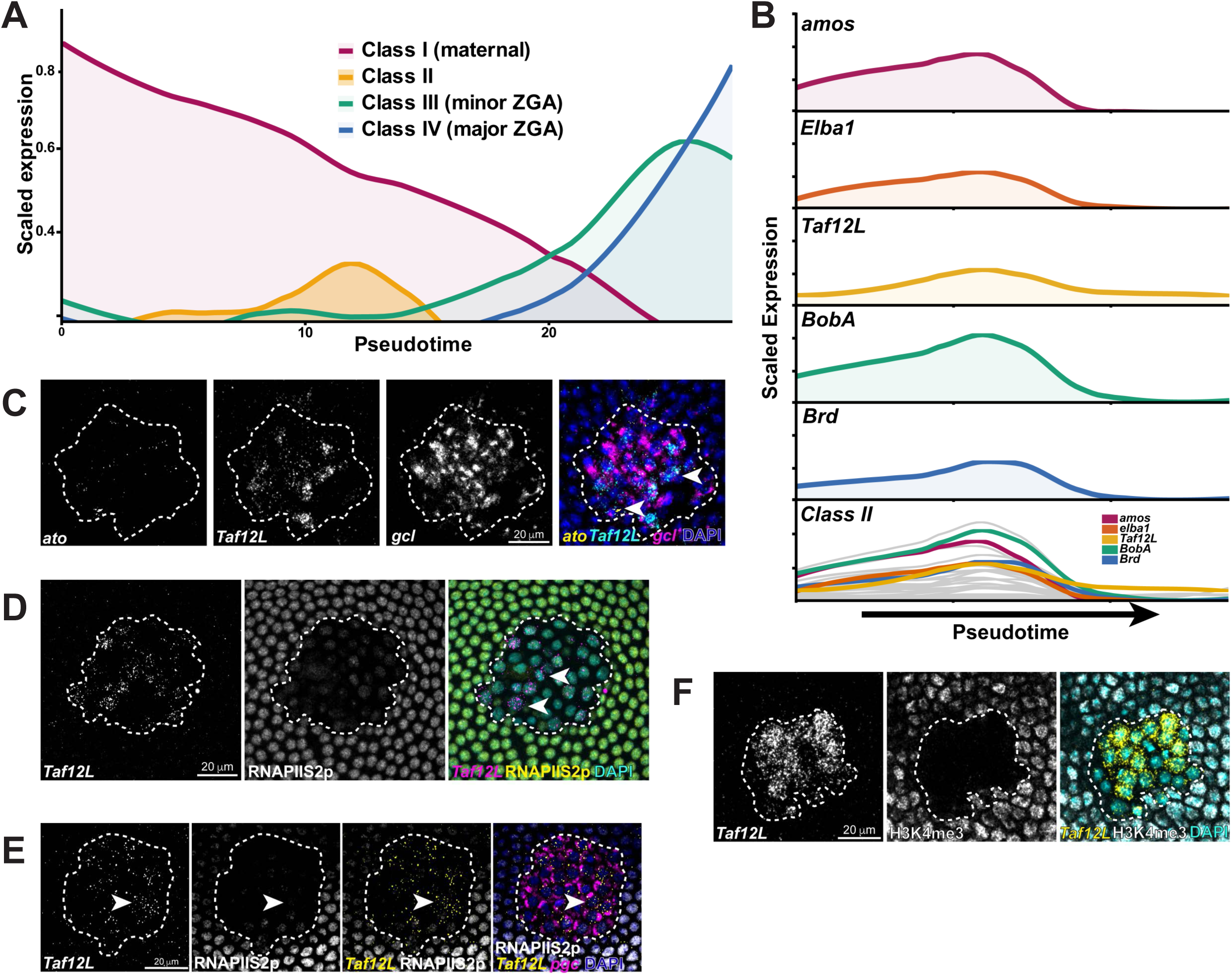
Class II genes are coordinately transcribed during germ cell specification. A. Scaled expression across pseudotime of the four gene classes: maternally deposited (Class I), early zygotic (Class II), minor ZGA (Class III), and major ZGA (Class IV). B. Scaled expression of five representative Class II genes, *amos*, *Elba1*, *Taf12L*, *BobA, and Brd* across pseudotime. The five representative genes are shown in color and the remaining Class II genes in grey. Expression is scaled from 0 to 1 for each gene. C. HCR *in situ* hybridization of Class II genes *ato* (yellow) and *Taf12L* (cyan) in stage 4 wild-type pole cells. Pole cells are marked by *gcl* (magenta) and outlined (dotted line). Nuclei are stained with DAPI (blue). Scale bar, 20 µm. D. Immunofluorescence for RNAPIIS2p (yellow) combined with HCR *in situ* hybridization of *Taf12L* (magenta) in wild-type embryos. Pole cells are outlined (dotted line). The surrounding soma displays high RNAPIIS2p, whereas pole cells show little to no signal, highlighting the low level of productive elongation in the germline. Arrowheads mark PGCs expressing *Taf12L* in the absence of RNAPIIS2p, demonstrating that Class II transcription occurs independently of productive RNAPII elongation. Scale bar, 20 µm. E. *Taf12L* is expressed in the absence of RNAPIIS2p and in the presence of *pgc* transcript. Immunofluorescence for RNAPIIS2p (white) combined with HCR *in situ* hybridization of *Taf12L* (yellow) and *pgc* (magenta). Pole cells outlined (dotted line). Arrowhead marks a transcribing PGC. Scale bar, 20 µm. F. Immunofluorescence for H3K4me3 (white) combined with HCR *in situ* hybridization of *Taf12L* (yellow). The surrounding soma displays strong H3K4me3, whereas pole cells show little to no signal, indicating that Class II transcription proceeds without this activating mark. Pole cells are outlined (dotted line). Nuclei stained with DAPI (cyan). Scale bar, 20 µm.

**fig. S3 (related to Figure 3):**
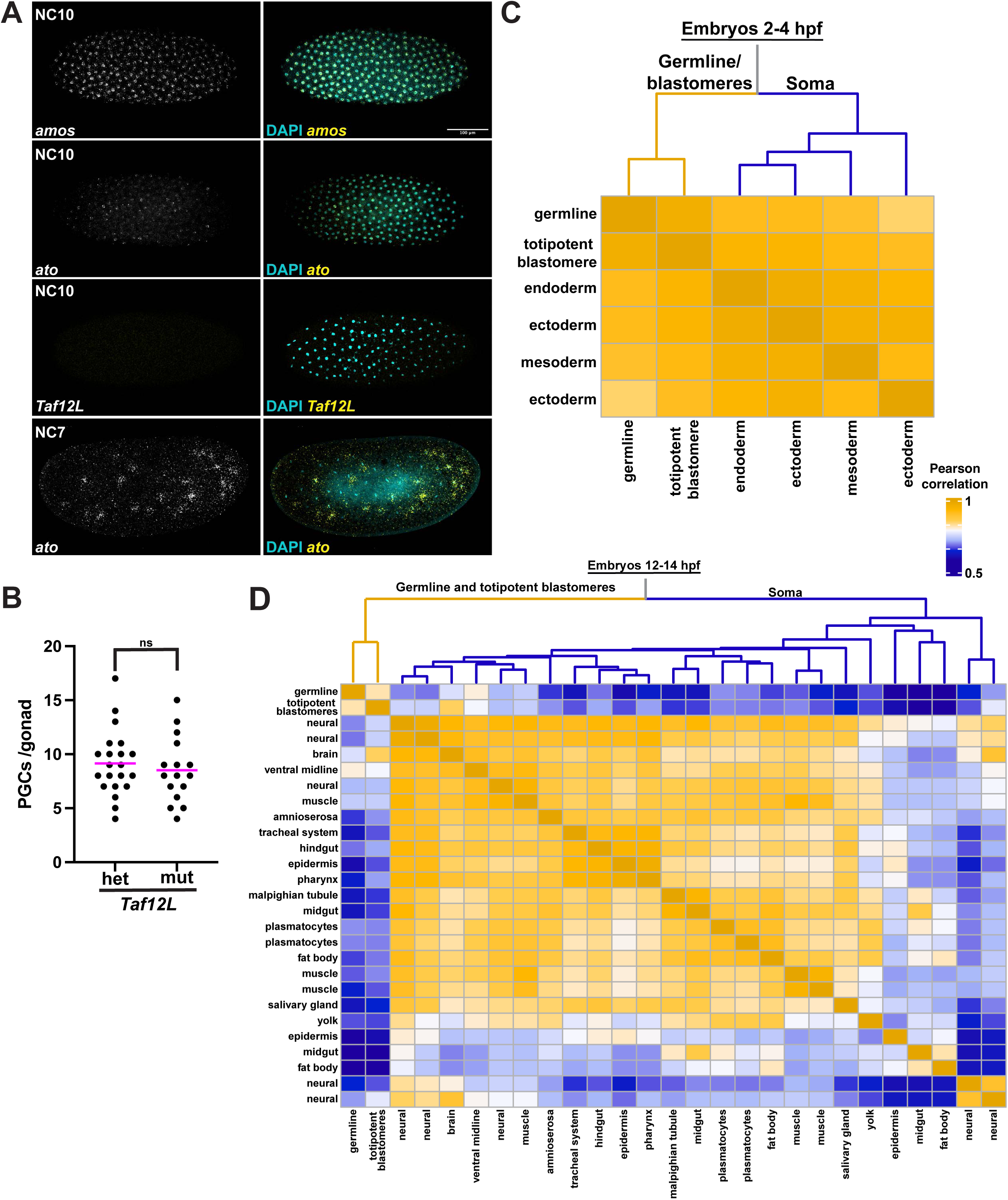
Germ cells and totipotent blastomeres share a chromatin state distinct from soma. A. HCR *in situ* hybridization of laterally mounted embryos showing only *amos*, *ato,* but not *Taf12L* expressed in the totipotent blastomeres at nuclear cycle 10. *Amos* and *ato* are expressed in totipotent blastomeres, while *Taf12L* is not. *ato* expression is also shown at the earliest time point at which expression is detected in the blastomeres, nuclear cycle 7. Nuclei stained with DAPI and shown in cyan. Scale bars, 100 µm. B. Number of PGCs that coalesced in the somatic gonad in control heterozygous and *Taf12L* mutant embryos. Each dot represents one gonad. Bar shows the mean. n = 21 and 16 gonads and 14 and 10 embryos, respectively; ns, not significant. C. Heatmap and dendrogram of pairwise Pearson correlation coefficients across snATAC-seq chromatin accessibility profiles of cell types from 2-4 hour embryos together with totipotent blastomeres from 0-2 hour embryos. At this early stage, the germline is most similar to the totipotent blastomeres, though all cell types are highly similar. snATAC-seq data from (*41*). D. Full dendrogram and heatmap of pairwise Pearson correlation coefficients from snATAC-seq chromatin accessibility profiles across all identified cell types including germ cells in embryos 12-14 hpf together with totipotent blastomeres from embryos 0-2 hpf. The germline and totipotent blastomeres form a distinct clade (yellow) from the differentiated somatic lineages (blue). snATAC-seq data from (*41*).

**fig. S4 (Related to Figure 4):**
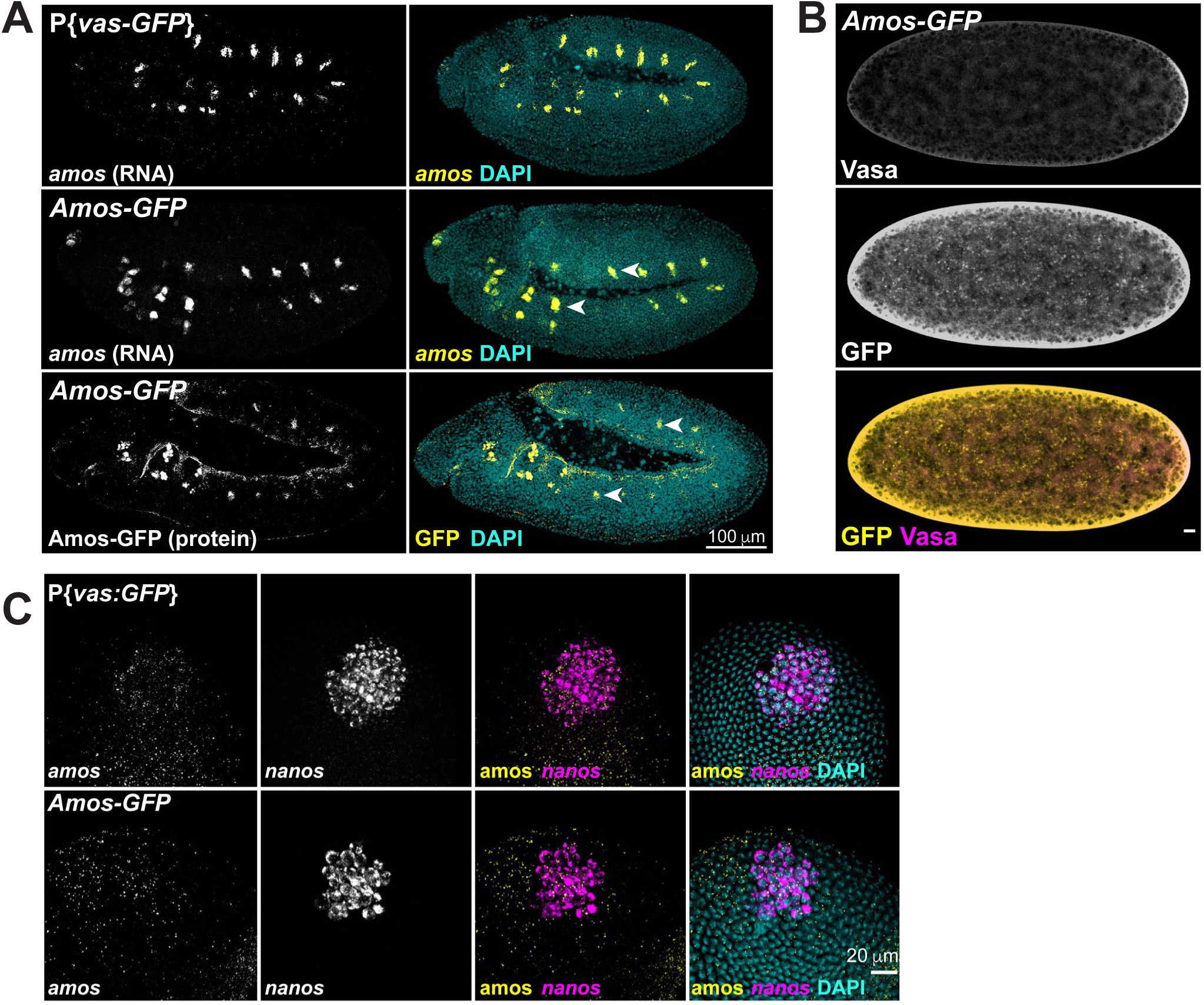
Validation of the Amos-GFP CRISPR line. A. Top: HCR *in situ* hybridization of *amos* RNA (yellow) in P{*vasa-GFP*} control embryos. Middle: HCR *in situ* hybridization of *amos* RNA (yellow) in Amos-GFP embryos. Bottom: immunofluorescence of GFP (yellow) in Amos-GFP embryos. *amos* RNA and Amos-GFP protein are each detected in neuronal precursor cells (marked by arrows), indicating that the GFP tag does not perturb *amos* expression in the soma. Nuclei shown in cyan. Scale bars, 100 µm. B. Immunofluorescence of Vasa protein (magenta) and Amos-GFP protein (yellow) in newly laid embryos. Broad, uniform GFP signal throughout the embryo indicates that Amos is maternally deposited as protein, with additional signal localized near nuclei. Scale bar, 20 µm. C. HCR *in situ* hybridization of *nanos* (magenta) and *amos* (yellow) in posterior-mounted embryos. *amos* expression is consistent between P{*vasa-GFP*} control and Amos-GFP lines. Pole cells outlined (dotted line). Nuclei stained with DAPI and shown in cyan. Scale bars, 20 µm.

**fig. S5 (Related to Figure 4):**
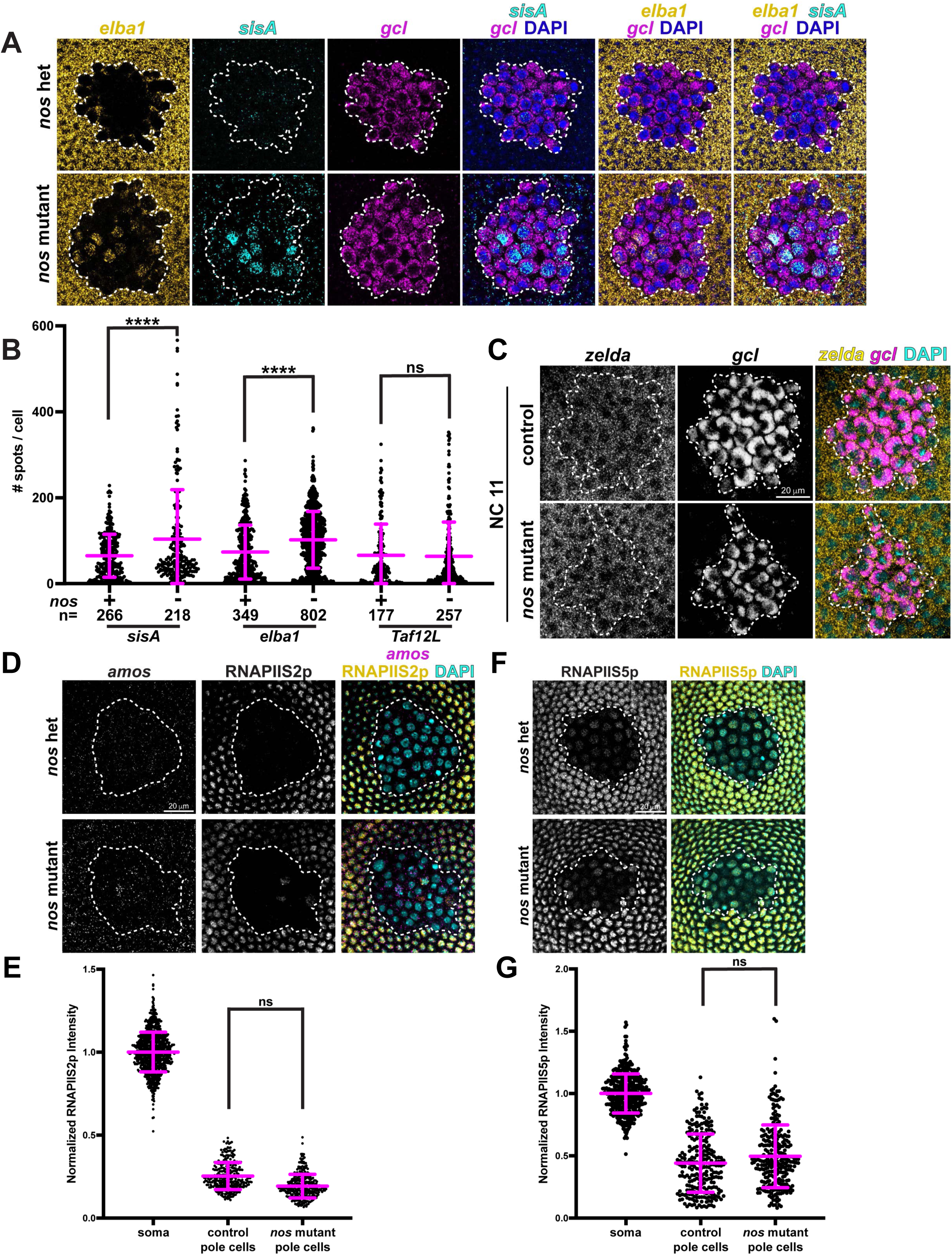
Nanos regulates Class II transcript levels in pole cells. A. *nanos* mutant pole cells accumulate elevated Class II transcripts. HCR *in situ* hybridization of control (*nanos* heterozygous) and *nanos* mutant embryos showing *elba1* (yellow, Class II gene), *sisA* (cyan, Class II gene), and *gcl* (magenta, germ cell marker). Many nuclei co-express *elba1* and *sisA*, while others express only one, consistent with the stochastic activation of Class II genes. Pole cells outlined (dotted line). Nuclei stained with DAPI (blue). Scale bar, 20 µm. B. Quantification of Class II signal in pole cells from embryos stained as in (A). Each dot represents one cell; bars indicate mean ± SD. Comparison between control and *nanos* mutant PGCs was performed using an unpaired Welch’s t-test; ****P < 0.0001. C. *zelda* RNA is uniformly present in pole cells at nuclear cycle 11. HCR *in situ* hybridization of control (*nanos* heterozygous) and *nanos* mutant embryos showing *zelda* (yellow) and *gcl* (magenta, germ cell marker). z*elda* RNA levels are similar between control and *nanos* mutant pole cells at this stage. Pole cells outlined (dotted line). Nuclei stained with DAPI (cyan). Scale bar, 20 µm. D. Immunofluorescence for RNAPIIS2p (yellow) combined with HCR *in situ* hybridization of *amos* RNA (magenta) in control (*nanos* heterozygous) and *nanos* mutant embryos. Germ cells outlined (dotted line). Nuclei stained with DAPI (cyan). Scale bar, 20 µm. E. Quantification of RNAPIIS2p signal in soma, control pole cells, and *nanos* mutant pole cells from embryos stained as in (D). Each dot represents one nucleus; bars indicate mean ± SD. Comparison between control and *nanos* mutant pole cells was performed on per-embryo means using a Wilcoxon rank-sum test (n = 17 and 15 embryos, respectively); ns, not significant. F. RNAPIIS5p phosphorylation is unchanged in *nanos* mutant embryos. Immunofluorescence for RNAPIIS5p (yellow) in control and *nanos* mutant embryos. Germ cells outlined (dotted line). Nuclei stained with DAPI (cyan). Scale bar, 20 µm. G. Quantification of RNAPIIS5p signal in soma, control pole cells, and *nanos* mutant pole cells from embryos stained as in (F). Each dot represents one nucleus; bars indicate mean ± SD. Comparison between control and *nanos* mutant pole cells was performed on per-embryo means using a Wilcoxon rank-sum test (n = 13 and 11 embryos, respectively); ns, not significant.

**fig. S6 (Related to Figure 5):**
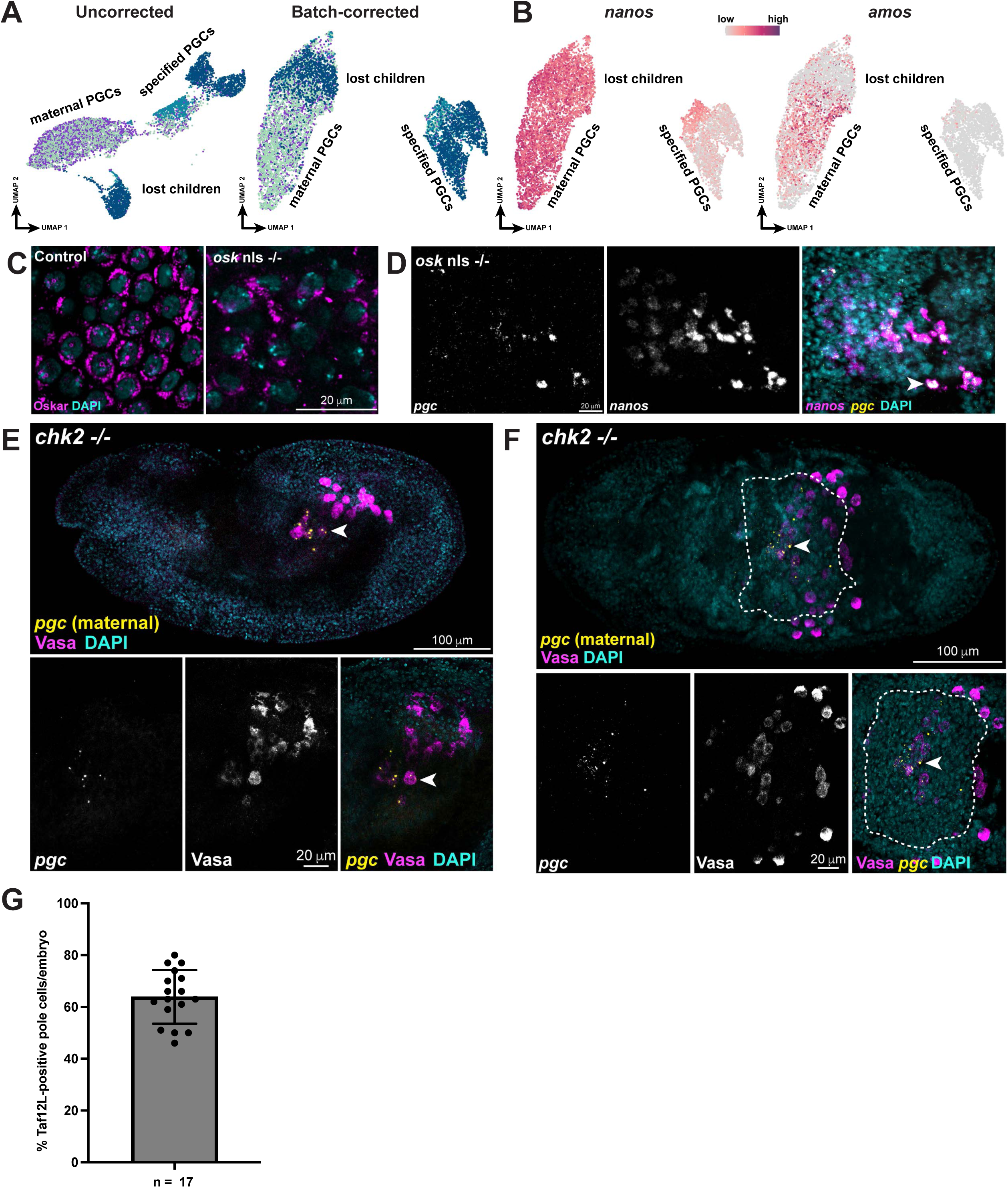
Cell death is a consequence, not the cause, of failed germ cell specification. A. Batch correction of scRNA-seq data using Harmony results in the lost children clustering within the early timepoints, indicating that their transcriptome resembles that of newly formed germ cells despite their late developmental stage. B. Feature plots of *nanos* and *amos* expression on the Harmony batch-corrected UMAP. Despite their late developmental age, the lost children nestle among the early timepoint germ cells, retaining the maternal transcript *nanos*. However, unlike the surrounding germ cells, the lost children do not express the Class II gene *amos*. C. Control embryos and embryos from *oskar* nuclear localization signal (NLS) mutant mothers stained for Oskar protein. *oskar* NLS mutants retain Oskar in the cytoplasm but lack nuclear Oskar granules, forming fewer pole cells. Scale bar, 20 µm. D. *oskar* NLS mutant embryos still produce lost children. HCR *in situ* hybridization of *oskar* NLS mutant embryos showing the maternal transcripts *pgc* (yellow) and *nanos* (magenta). Pole cells outlined (dotted line). Nuclei stained with DAPI (cyan). Arrowhead marks lost children. Scale bars, 20 µm. E. Rescuing cell death does not rescue the lost children. HCR *in situ* hybridization and immunofluorescence of stage 11 *chk2* mutant (maternal and zygotic) embryo (top) with high-magnification imaging of germ cells (bottom). *pgc* (yellow) marks lost children; Vasa protein (magenta) is enriched in healthy PGCs. Nuclei stained with DAPI (cyan). Scale bars, 100 µm and 20 µm. **(E)** Lateral view, arrowheads mark lost children. F. Same as E but ventral view, lost children shown within the gut (dashed line) based on endoderm marker expression. G. Quantification of the percentage of *Taf12L*-positive pole cells per wild-type embryo. Each dot represents one embryo (n = 17); bar shows the mean number of pole cells (63.88), error bars represent standard deviation. The ∼40% of pole cells that do not express *Taf12L* corresponds to the proportion of germ cells that become lost children.

**Additional Data Table S1:** Genes differentially expressed between XY and XX germ cells Differentially expressed genes distinguishing XY and XX PGC clusters, identified using Seurat’s FindMarkers (Wilcoxon rank-sum test). For each gene: gene symbol; p-value; average log2 fold-change (XY relative to XX); the fraction of cells expressing the gene in the XY (pct.1, cluster 5) and XX (pct.2, cluster 6) clusters; adjusted p-value (Bonferroni-corrected); and chromosomal location. Of these markers, 99 are X-linked and show lower expression in XY than XX, reflecting the 2-fold higher expression of X-linked genes in XX PGCs in the absence of dosage compensation. The single non-X marker is *FDY*, a Y-linked gene expressed specifically in XY PGCs.

**Additional Data Table S2:** Class II genes: transcript features, regulation, and function Sheet 1: Class II genes. For each gene: gene symbol; FlyBase gene identifier (FBID); annotated transcript(s); the full transcriptional unit length (nucleotide (nt), genomic span from the 5’ to 3’ end, including introns); the processed mRNA length (nt, exons only); intron count and intron sizes; whether the gene is transcribed in the totipotent blastomeres of the early embryo during nuclear cycles 7-13 (*29*); its RNA polymerase II pausing status in blastomeres (*28*); whether it is a Zelda target based on Zelda-dependent expression (*34, 35*); whether its promoter is bound by Zelda (*33*); and whether it is a Nanos regulatory target (“yes” = significantly targeted, “yes-partial” = reduced but not significantly, “no” = not targeted, “not detected” = not detected in the dataset) (*49*). Note: lncRNA:CR43264 shows Class II-like expression in our atlas but was excluded from all other analyses because it is a long non-coding RNA rather than a protein-coding gene. Sheet 2: Class II function. UniProt function is provided for each Class II gene (data from FlyBase). Functional category was assigned based on the UniProt function and available literature.

## References

1. R. Lehmann, Germline stem cells: origin and destiny. Cell stem cell 10, 729–739 (2012).

2. C. G. Extavour, M. Akam, Mechanisms of germ cell specification across the metazoans: epigenesis and preformation. Development 130, 5869–5884 (2003).

3. M. Hemberger, W. Dean, W. Reik, Epigenetic dynamics of stem cells and cell lineage commitment: digging Waddington’s canal. Nature reviews. Molecular cell biology 10, 526–537 (2009).

4. T. Chen, S. Y. Dent, Chromatin modifiers and remodellers: regulators of cellular differentiation. Nat Rev Genet 15, 93–106 (2014).

5. R. Chen, S. Grill, B. Lin, M. Saiduddin, R. Lehmann, Origin and establishment of the germline in Drosophila melanogaster. Genetics 229, (2025).

6. R. M. Cinalli, P. Rangan, R. Lehmann, Germ cells are forever. Cell 132, 559–562 (2008).

7. T. Graf, T. Enver, Forcing cells to change lineages. Nature 462, 587–594 (2009).

8. A. Nakamura, G. Seydoux, Less is more: specification of the germline by transcriptional repression. Development 135, 3817–3827 (2008).

9. N. U. Siddiqui et al., Genome-wide analysis of the maternal-to-zygotic transition in Drosophila primordial germ cells. Genome Biol 13, R11 (2012).

10. N. L. Vastenhouw, W. X. Cao, H. D. Lipshitz, The maternal-to-zygotic transition revisited. Development 146, (2019).

11. M. Poirie, E. Niederer, M. Steinmann-Zwicky, A sex-specific number of germ cells in embryonic gonads of Drosophila. Development 121, 1867–1873 (1995).

12. G. Weidinger et al., dead end, a novel vertebrate germ plasm component, is required for zebrafish primordial germ cell migration and survival. Curr Biol 13, 1429–1434 (2003).

13. J. Stallock, K. Molyneaux, K. Schaible, C. M. Knudson, C. Wylie, The pro-apoptotic gene Bax is required for the death of ectopic primordial germ cells during their migration in the mouse embryo. Development 130, 6589–6597 (2003).

14. B. E. Richardson, R. Lehmann, Mechanisms guiding primordial germ cell migration: strategies from different organisms. Nature reviews. Molecular cell biology 11, 37–49 (2010).

15. M. Slaidina, R. Lehmann, Quantitative Differences in a Single Maternal Factor Determine Survival Probabilities among Drosophila Germ Cells. Curr Biol 27, 291–297 (2017).

16. A. D. Renault, vasa is expressed in somatic cells of the embryonic gonad in a sex-specific manner in Drosophila melanogaster. Biol Open 1, 1043–1048 (2012).

17. K. Hanyu-Nakamura, H. Sonobe-Nojima, A. Tanigawa, P. Lasko, A. Nakamura, Drosophila Pgc protein inhibits P-TEFb recruitment to chromatin in primordial germ cells. Nature 451, 730–733 (2008).

18. R. G. Martinho, P. S. Kunwar, J. Casanova, R. Lehmann, A noncoding RNA is required for the repression of RNApolII-dependent transcription in primordial germ cells. Curr Biol 14, 159–165 (2004).

19. S. Grill, A. Riley, M. Selvaraj, R. Lehmann, HP6/Umbrea is dispensable for viability and fertility, suggesting essentiality of newly evolved genes is rare. Proc Natl Acad Sci U S A 120, e2309478120 (2023).

20. C. Rushlow, H. Doyle, T. Hoey, M. Levine, Molecular characterization of the zerknullt region of the Antennapedia gene complex in Drosophila. Genes Dev 1, 1268–1279 (1987).

21. C. E. Eichler, A. C. Hakes, B. Hull, E. R. Gavis, Compartmentalized oskar degradation in the germ plasm safeguards germline development. Elife 9, (2020).

22. M. Van Doren, A. L. Williamson, R. Lehmann, Regulation of zygotic gene expression in Drosophila primordial germ cells. Curr Biol 8, 243–246 (1998).

23. Y. R. Li et al., Trajectory mapping of the early. Genome Res 31, 1011–1023 (2021).

24. R. Ota, M. Hayashi, S. Morita, H. Miura, S. Kobayashi, Absence of X-chromosome dosage compensation in the primordial germ cells of Drosophila embryos. Sci Rep 11, 4890 (2021).

25. D. Peng et al., Organogenetic transcriptomes of the Drosophila embryo at single cell resolution. Development 151, (2024).

26. G. Seydoux, M. A. Dunn, Transcriptionally repressed germ cells lack a subpopulation of phosphorylated RNA polymerase II in early embryos of Caenorhabditis elegans and Drosophila melanogaster. Development 124, 2191–2201 (1997).

27. W. Tadros, H. D. Lipshitz, The maternal-to-zygotic transition: a play in two acts. Development 136, 3033–3042 (2009).

28. K. Chen et al., A global change in RNA polymerase II pausing during the Drosophila midblastula transition. Elife 2, e00861 (2013).

29. J. C. Kwasnieski, T. L. Orr-Weaver, D. P. Bartel, Early genome activation in Drosophila is extensive with an initial tendency for aborted transcripts and retained introns. Genome Res 29, 1188–1197 (2019).

30. S. De Renzis, O. Elemento, S. Tavazoie, E. F. Wieschaus, Unmasking activation of the zygotic genome using chromosomal deletions in the Drosophila embryo. PLoS Biol 5, e117 (2007).

31. M. Zalokar, Transplantation of nucle in Drosophila melanogaster. Proc Natl Acad Sci U S A 68, 1539–1541 (1971).

32. E. Posfai et al., Evaluating totipotency using criteria of increasing stringency. Nat Cell Biol 23, 49–60 (2021).

33. M. M. Harrison, X. Y. Li, T. Kaplan, M. R. Botchan, M. B. Eisen, Zelda binding in the early Drosophila melanogaster embryo marks regions subsequently activated at the maternal-to-zygotic transition. PLoS Genet 7, e1002266 (2011).

34. C. Y. Nien et al., Temporal coordination of gene networks by Zelda in the early Drosophila embryo. PLoS Genet 7, e1002339 (2011).

35. H. L. Liang et al., The zinc-finger protein Zelda is a key activator of the early zygotic genome in Drosophila. Nature 456, 400–403 (2008).

36. M. L. Huang, C. H. Hsu, C. T. Chien, The proneural gene amos promotes multiple dendritic neuron formation in the Drosophila peripheral nervous system. Neuron 25, 57–67 (2000).

37. A. P. Jarman, Y. Grau, L. Y. Jan, Y. N. Jan, atonal is a proneural gene that directs chordotonal organ formation in the Drosophila peripheral nervous system. Cell 73, 1307–1321 (1993).

38. A. P. Jarman, E. H. Grell, L. Ackerman, L. Y. Jan, Y. N. Jan, Atonal is the proneural gene for Drosophila photoreceptors. Nature 369, 398–400 (1994).

39. X. Chen, M. Hiller, Y. Sancak, M. T. Fuller, Tissue-specific TAFs counteract Polycomb to turn on terminal differentiation. Science 310, 869–872 (2005).

40. M. Hiller et al., Testis-specific TAF homologs collaborate to control a tissue-specific transcription program. Development 131, 5297–5308 (2004).

41. D. Calderon et al., The continuum of Drosophila embryonic development at single-cell resolution. Science 377, eabn5800 (2022).

42. A. Gaspar-Maia, A. Alajem, E. Meshorer, M. Ramalho-Santos, Open chromatin in pluripotency and reprogramming. Nature reviews. Molecular cell biology 12, 36–47 (2011).

43. E. Meshorer, T. Misteli, Chromatin in pluripotent embryonic stem cells and differentiation. Nature reviews. Molecular cell biology 7, 540–546 (2006).

44. R. Lehmann, C. Nusslein-Volhard, The maternal gene nanos has a central role in posterior pattern formation of the Drosophila embryo. Development 112, 679–691 (1991).

45. C. Wang, R. Lehmann, Nanos is the localized posterior determinant in Drosophila. Cell 66, 637–647 (1991).

46. S. Kobayashi, M. Yamada, M. Asaoka, T. Kitamura, Essential role of the posterior morphogen nanos for germline development in Drosophila. Nature 380, 708–711 (1996).

47. C. Wreden, A. C. Verrotti, J. A. Schisa, M. E. Lieberfarb, S. Strickland, Nanos and pumilio establish embryonic polarity in Drosophila by promoting posterior deadenylation of hunchback mRNA. Development 124, 3015–3023 (1997).

48. T. Raisch et al., Distinct modes of recruitment of the CCR4-NOT complex by Drosophila and vertebrate Nanos. EMBO J 35, 974–990 (2016).

49. M. Marhabaie, T. H. Wharton, S. Y. Kim, R. P. Wharton, Widespread regulation of the maternal transcriptome by Nanos in Drosophila. PLoS Biol 22, e3002840 (2024).

50. K. Ikenishi, T. Ohno, T. Komiya, Ectopic germline cells in embryos of Xenopus laevis. Dev Growth Differ 49, 561–570 (2007).

51. K. E. Kistler et al., Phase transitioned nuclear Oskar promotes cell division of. Elife 7, (2018).

52. S. Strome, R. Lehmann, Germ versus soma decisions: lessons from flies and worms. Science 316, 392–393 (2007).

53. Y. Hayashi, M. Hayashi, S. Kobayashi, Nanos suppresses somatic cell fate in Drosophila germ line. Proc Natl Acad Sci U S A 101, 10338–10342 (2004).

54. D. C. Hamm et al., A conserved maternal-specific repressive domain in Zelda revealed by Cas9-mediated mutagenesis in. Plos Genetics 13, (2017).

55. W. A. Pastor et al., TFAP2C regulates transcription in human naive pluripotency by opening enhancers. Nat Cell Biol 20, 553–564 (2018).

56. N. Grabole et al., Prdm14 promotes germline fate and naive pluripotency by repressing FGF signalling and DNA methylation. EMBO Rep 14, 629–637 (2013).

57. S. Weber et al., Critical function of AP-2 gamma/TCFAP2C in mouse embryonic germ cell maintenance. Biol Reprod 82, 214–223 (2010).

58. M. Yamaji et al., Critical function of Prdm14 for the establishment of the germ cell lineage in mice. Nat Genet 40, 1016–1022 (2008).

59. E. Magnúsdóttir et al., A tripartite transcription factor network regulates primordial germ cell specification in mice. Nat Cell Biol 15, 905–915 (2013).

60. M. Saitou, M. Yamaji, Primordial germ cells in mice. Cold Spring Harb Perspect Biol 4, (2012).

61. K. Hayashi, H. Ohta, K. Kurimoto, S. Aramaki, M. Saitou, Reconstitution of the mouse germ cell specification pathway in culture by pluripotent stem cells. Cell 146, 519–532 (2011).

62. I. Korsunsky et al., Fast, sensitive and accurate integration of single-cell data with Harmony. Nat Methods 16, 1289–1296 (2019).

63. E. P. Consortium, An integrated encyclopedia of DNA elements in the human genome. Nature 489, 57–74 (2012).

64. M. Pachitariu, C. Stringer, Cellpose 2.0: how to train your own model. Nat Methods 19, 1634–1641 (2022).

65. C. Stringer, T. Wang, M. Michaelos, M. Pachitariu, Cellpose: a generalist algorithm for cellular segmentation. Nat Methods 18, 100–106 (2021).

66. C. P. Stringer, M., Cellpose-SAM: superhuman generalization for cellular segmentation. BioRxiv, (2025).

67. A. H. Albert Dominguez Mantes, Irina Khven, Anjalie Schlaeppi, Eftychia Kyriacou, Georgios Tsissios, Can Aztekin, Joachim Lingner, Gioele La Manno, Martin Weigert, Spotiflow: accurate and efficient spot detection for imaging-based spatial transcriptomics. BioRxiv, (2024).

68. M. Starz-Gaiano, R. Lehmann, Moving towards the next generation. Mech Dev 105, 5–18 (2001).

